# Mutant evolution in spatially structured and fragmented populations

**DOI:** 10.1101/817387

**Authors:** Dominik Wodarz, Natalia L Komarova

## Abstract

Understanding mutant evolution in spatially structured systems is crucially important for a range of biological systems, including bacterial populations and cancer. While previous work has shown that the mutation load is higher in spatially structured compared to well-mixed systems for neutral mutants in the absence of cell death, we demonstrate a significantly higher degree of complexity, using a comprehensive computational modeling approach that takes into account different mutant fitness, cell death, and different population structures. While an agent-based model assuming nearest neighbor interactions predicts a higher abundance of neutral or advantageous mutants compared to well-mixed systems of the same size, we show that for disadvantageous mutants, results depend on the nature of the disadvantage. In particular, if the disadvantage occurs through higher death rates, as opposed to lower reproduction rates, the result is the opposite, and a lower mutation load occurs in spatial compared to mixed systems. Interestingly, we show that in all cases, the same results are observed in fragmented patch models, where individuals can migrate to randomly chosen patches, thus lacking a strict spatial component. Hence, the results reported for spatial models are the consequence of population fragmentation, and not spatial restrictions per se. We further derive growth laws that characterize the expansion of wild type and mutant populations in different dimensionalities. For example, we find that while disadvantageous mutant abundance scales with the total population size, N, neutral mutants grow faster (as N^2^ for 1D, as N^3/2^ in 2D, and as N^4/3^ in 3D). Advantageous mutants scale with the cube of N in 1D and with the square of N in higher dimensions. These laws are universal (as long as mutants remain a minority) and independent of “microscopic” modeling details.

## Introduction

The dynamics of mutant creation and invasion are relatively well understood under a variety of conditions and assumptions, mostly assuming perfect mixing of individuals. In the context of constant populations, the fixation probability of mutants as well as fixation times have been thoroughly defined under various assumptions in the population genetics literature [1,2]. The emergence of mutants in exponentially growing bacterial populations is also well studied, based on the famous Luria-Delbruck experiments [3] and the resulting rich theoretical framework [4-7]. This has been instrumental for understanding the principles according to which antibiotic-resistant microbes emerge [8], and has also been applied to studying the emergence of drug resistance in some cancers [9-11]. The majority of tumors, however, are characterized by 2D and 3D spatial structures, and so is the growth of bacteria in biofilms. Recent experimental and theoretical work [12] has extended our understanding of mutant emergence to spatially structured populations. An excess of mutational jackpot events was observed in spatial compared to well-mixed systems. Such events result from mutations arising at the surface of expanding, spatially structured populations, surfing at the edge of range expansions, and appearing as mutant “sectors” or “slices”. These jackpot events can occur relatively late in the expansion process, which is in contrast to well-mixed systems in which mutational jackpot events can only occur early on in the population growth process [12]. Hence, overall, the average number of mutants when the total population reaches a given threshold size is significantly larger in spatial compared to non-spatial settings [12]. This work was done under the assumption that cells do not die, and theory and computations were mostly developed in the context of neutral mutants. Computational analysis was performed using an agent-based modeling approach.

Here, we expand this work and investigate the dynamics of mutant emergence and growth in spatially structured cell populations assuming varying death rates, different mutant fitness, different dimensionalities of space, and different spatial modeling approaches. One of the two main messages of this paper is to report interesting dynamics observed for disadvantageous mutants, which could apply for example to drug-resistant mutants that emerge before the onset of therapy. If the disadvantage is caused by a larger death rate of the mutant cells, then we find that in contrast to other scenarios, the number of mutants at a given size is larger in a well-mixed compared to the spatial system. If, on the other hand, the fitness disadvantage arises because of a slower replication rate, then more mutants are found in the spatial compared to the non-spatial system, similar to the results obtained for neutral or advantageous mutants. We further find identical results in a patch model, where local within-patch dynamics are governed by perfect mixing, but individuals migrate to other patches. Interestingly, the results do not depend on the assumption that patches are spatially arranged, with migration of individuals to nearest neighboring patches. The same outcomes are observed if migration can occur to any randomly chosen patch in the system. Therefore, the properties of mutant growth in the spatial agent-based model might not be the direct consequence of spatial dynamics, but the consequence of population fragmentation.

The second focus of this paper is to derive universal growth laws that govern cell expansion in spatially constrained models. The so-called “surface growth” law of homogeneous cell colonies in space has previously been described in experiments [13-15] and in the modeling literature [16-21]. Here we study the laws of mutant generation, spread, and competition with the wild type individuals, in the context of spatially restricted colony expansion. We derive formulas that relate the expected number of disadvantageous, neutral, and advantageous mutants to the total population size in different spatial dimensions.

## Generation and spread of mutants in spatial and non-spatial models

We used a 2-dimensional agent-based model and a patch model (see Materials and Methods) to explore the spread of mutants in spatial and non-spatial growth processes. Denote by L_w_ and L_m_ the division rates of wild type and mutant cells, and by D_w_ and D_m_ their respective death rates. Below we report the results for neutral, disadvantageous, and advantageous mutants.

### Neutral mutants

First we used the 2-dimensional agent-based model under the same assumptions as used in [12], i.e. with neutral mutants and zero death rates (L_w_=L_m_>0, D_w_=D_m_=0). The same type of dynamics are observed as previously reported, with mutant clones either being engulfed by wild-type cells after creation, or mutant clones establishing growing sectors. The average number of mutants at size M is significantly larger for the spatial compared to the non-spatial system (not shown).

Similar results are observed under the assumption that cells can die (L_w_=L_m_>0, D_w_=D_m_>0). Mutants either grow as expanding sectors or are engulfed by the wild-type cells after temporary expansion (Figure 1, inset in ai). The number of mutants at population size M is always larger in the spatial compared to the non-spatial system (Figure 1a). The extent of the difference is larger for higher death rates and lower reproduction rates, i.e. for slower growing population (Figure 1a).

**Fig 1.**
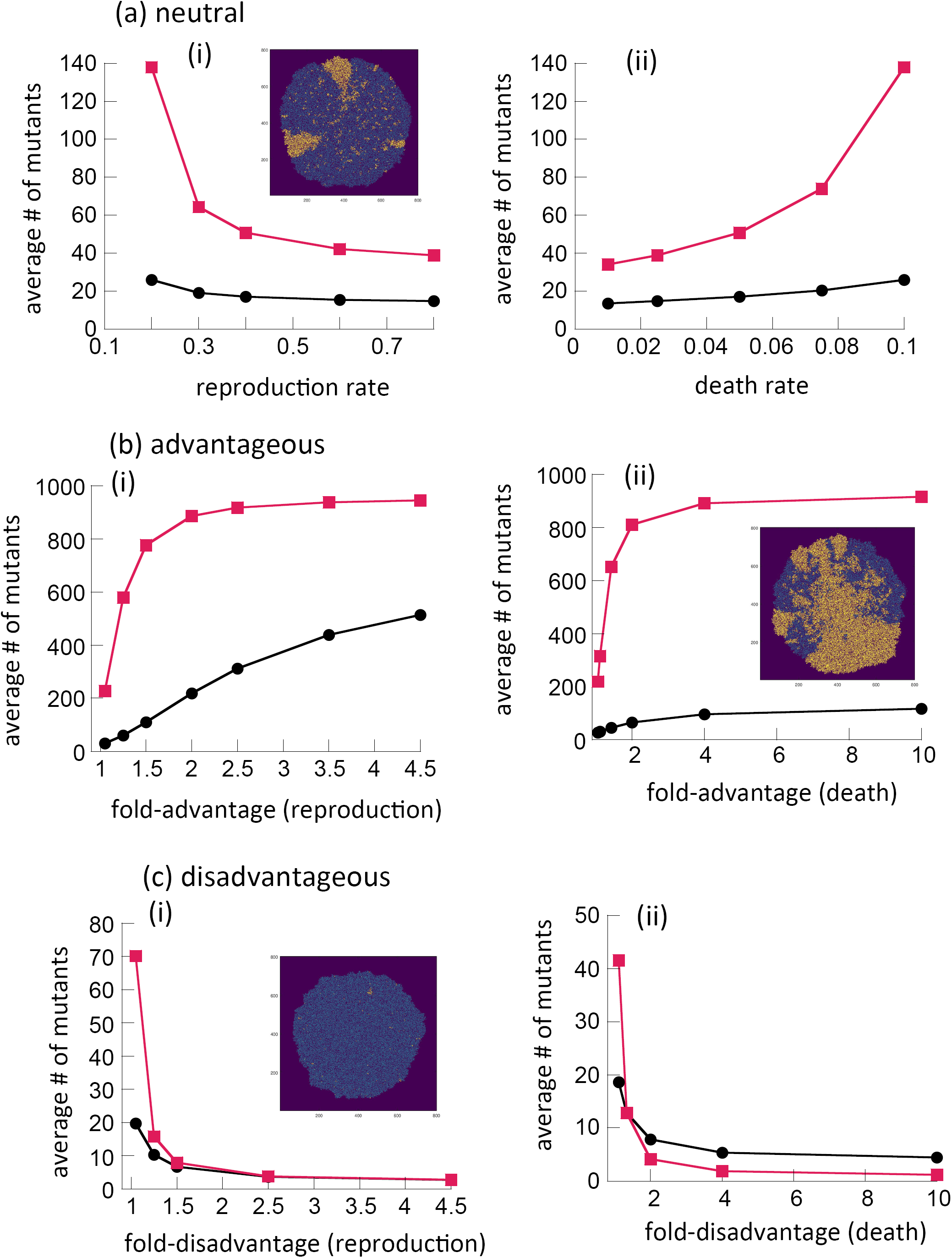
Comparison of the number of mutants in 2D spatial agent-based model simulations (red) and a well-mixed system (black). The lines represent the mean numbers of mutants in the spatial and non-spatial systems at equal size, N=10^3^. (a) Neutral mutants, as a function of the division rate (i) and the death rate (ii). (b) Advantageous mutants, characterized by increased division rates (bi) and decreased death rates (ii), as a function of the fold-advantage, (c) Disdvantageous mutants characterized by decreased division rates (ci) and increased death rates (cii), as a function of the fold-disadvantage. Typical 2D spatial agent-based simulations of range expansion dynamics are shown in the insets for each mutant type. The parameters (unless otherwise indicated in figure) are: L=0.2, D=0.1, u=2×10^−3^. For each parameter combination, from 2×10^6^ to 3×10^7^ repeats were performed; shown are the means; standard errors are too small to see.

Results of this comparison were qualitatively similar to those obtained from the patch model. The number of neutral mutants at population size M (assumed much smaller than the maximum system size) was always higher for the spatial (patch) compared to the well-mixed system (Fig S2(A) of the Supplement). Interestingly, this result holds for different spatial organizations of the patch model. In the most spatially restricted system, individuals can only migrate to and from the eight nearest neighboring patches. In an alternative model, migration is allowed between any patches regardless of their location. In either case, a patch model produces significantly more mutants than the mass-action system. This suggests that it is not the spatial arrangement per se but fragmentation of the system that may be responsible for the observed increased number of mutants. The difference is more pronounced for larger cell death rates (Fig S2(A) of the Supplement).

### Advantageous mutants

If the mutant is advantageous, the dynamics are similar as those observed for neutral mutants. First we assume that the advantage is given by a larger division rate of the mutant cells. The number of mutants at population size M is always larger in the spatial compared to the well-mixed setting. This difference tends to become larger for more pronounced fitness advantages (Figure 1bi). In addition, higher death rates lead to a larger difference between the number of mutants in spatial and non-spatial settings (Fig S1(a) in the Supplement). Mutants can again either grow as expanding sectors, or show a temporary growth phase before being engulfed by wild-type cells, see inset in Fig 1bii. Similar results are obtained if we assume that the mutant advantage is given through a reduced death rate of mutant cells (Figure 1bii and Fig S1(b) in the Supplement). These conclusions remain robust if we use the patch model (either spatially constrained or with random migration between any two patches) instead of the agent-based model (Fig S2(B) of the Supplement).

### Disadvantageous mutants

Disadvantageous mutants are very unlikely to grow as sustained sectors, especially if the disadvantage is more pronounced (inset in Figure 1ci). In the absence of death, after creation, mutants undergo a few cell divisions and are then engulfed by the expanding wild-type cell population; in the presence of cell death, they form mutant “islands”, which can become repeatedly generated by mutations and tend to be outcompeted by wild-type cells.

The average number of mutants when the overall population reaches size M depends on spatial structure in a more complex way, compared to the case of neutral mutants. First, we assume that the fitness difference lies in the division rate of the cells (figure 1ci). In this case, we observe that the average number of mutants is always larger for the spatial compared to the well-mixed simulations. The extent of the difference, however, becomes very small as the extent of the disadvantage grows (Figure 1ci). Hence, unless the mutant is almost neutral, the increase in the number of mutants in the spatial compared to the non-spatial system becomes negligible. In addition, the difference is most pronounced for small death rates and diminishes for larger death rates (Fig S1(c) of the Supplement).

A different result is observed if the lower fitness of the mutant strain is brought about by a higher death rate of mutant cells. If the difference in death rates lies above a threshold level, the average number of mutants at size M is observed to be larger in well-mixed compared to spatial simulations (Fig 1cii), which is the opposite trend compared to the previous cases, and also the opposite result compared to those reported in [12]. The lower the difference in death rates, the less pronounced this effect. If the difference in death rates between mutant and wild-type cells falls below a threshold, then the results reverse and become similar to those obtained for neutral mutants: the average number of mutants becomes larger in spatial compared to well-mixed simulations (Figure 1cii). Lower reproduction rates result in more pronounced differences between the number of mutants in spatial and non-spatial settings (Fig S1(d) of the Supplement). An analysis of the spatial stochastic model is developed in Section 3 of the Supplement. Using the pair approximation, we derive an approximation for the selection-mutation balance of mutants away from the colony boundary (formula (36)). This theory predicts patterns similar to those described above.

To confirm the robustness of these results, we performed simulations with disadvantageous mutants in a patch model (as before, two migration schemes were used: to nearest patches and to all patches). Again, the outcome of the dynamics depends on the parameters upon which the disadvantage is based, see Fig 2. If the mutant has a lower division rate than the wild-type, the number of mutants at population size M is larger for the spatial than for the well-mixed scenario. This difference is largest if cells do not die, and diminishes with increasing cell death rates. If, however, the mutant is characterized by a larger death rate than the wild-type, then the opposite result is obtained: the number of mutants at population size M is smaller in the spatial than in the well-mixed system (as long as the difference in the death rate lies above a threshold). Again, the results are qualitatively similar for the spatially restricted (nearest neighbor) and non-restricted (migration to all patches) models (see yellow and green lines in the graph of Fig 2).

**Fig. 2.**
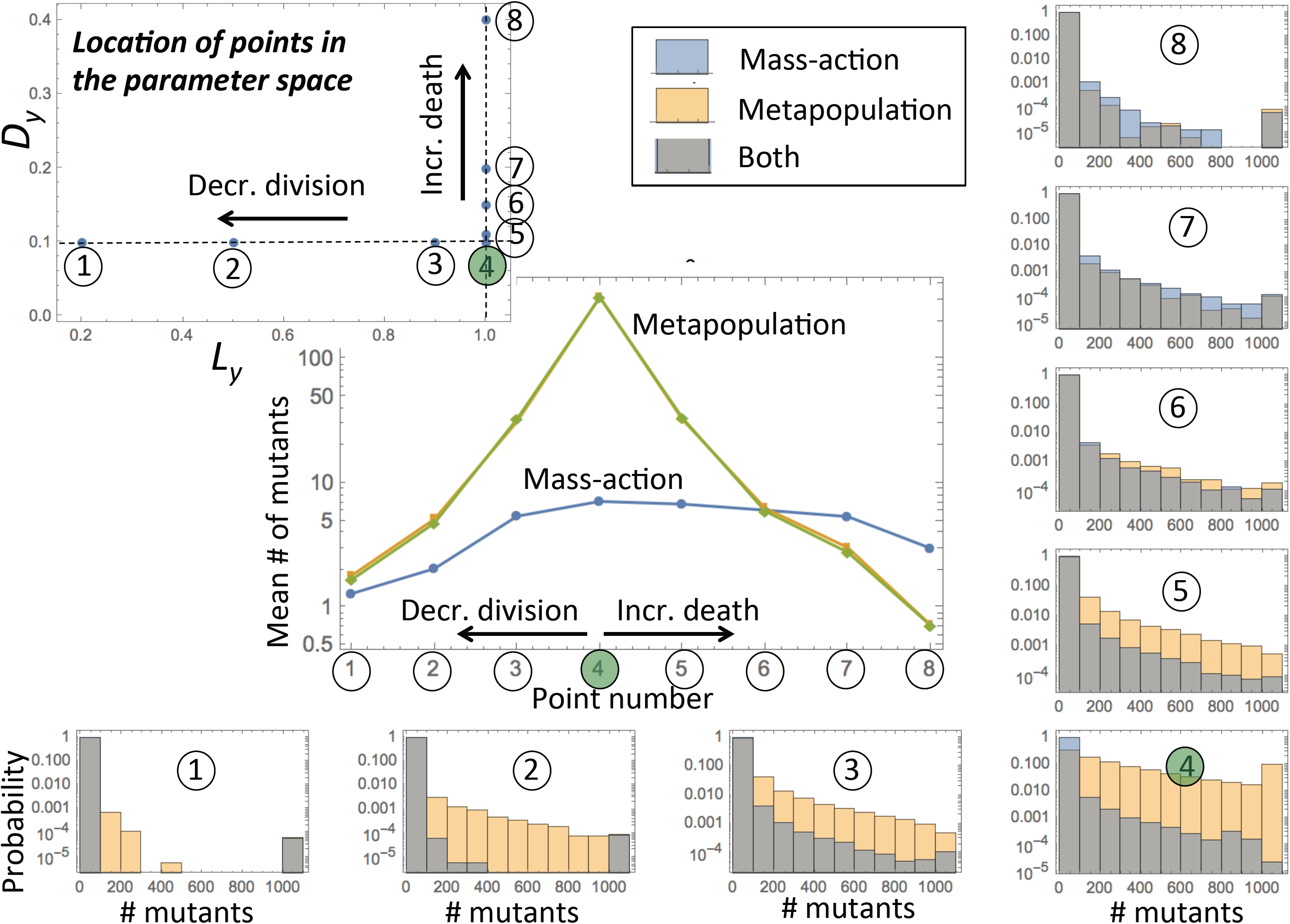
A systematic study of the number of disadvantageous mutants in mass-action (blue bars in histograms) and in patch model (yellow bars). Data are presented for 8 parameter combinations: 1-3 with mutants of decreased division rates, 4 (the green point) with neutral mutants, and 5-8 with mutants of increased death rate. We observe that the number of mutants in the patch models (with nearest neighbor migrations and global migrations, depicted by yellow and green lines in the graph) becomes smaller than that in the mass-action model (blue line) if the disadvantage through death is sufficiently large. The numerical probability distributions for the numbers of mutants are also presented for well-mixed and patch (with nearest neighbor migration) models. The rest of the parameters are as follows: u=10^−3^, µ=10^−5^, k=100, 100 patches; the number of mutants evaluated at total population 10^3^.

Deterministic (ODE) versions of the patch models are developed and analyzed in Section 2 of the Supplement. In particular, Section 2.5 of the Supplement provides approximate formulas for the numbers of mutants in a metapopulation model and shows for what division and death parameters the number of disadvantageous mutants is higher (lower) in the deterministic metapopulation model compared to the mass-action model. One of the consequences is that for mutants with lower division rates, more mutants occur in a deterministic metapopulation model, and for mutants with sufficiently high death rates, there are more mutants in mas-action.

### Disadvantageous mutants: an intuitive explanation of growth patterns

An intuitive explanation of this phenomenon can be built by observing the growth patterns of disadvantageous mutants in a single patch, starting from a single wild type cell (see Sections 2.5-2.6 of the Supplement for details). Typically, as the total population increases and reaches its carrying capacity, the mean number of mutants is an increasing function of the total population size (which eventually on average saturates at the selection mutation balance). The number of mutants however does not grow proportional to the total population size, in fact, in some cases the percentage of mutants increases with size, and in others it decreases with size. It turns out that mutants characterized by decreased division rates, which grow relatively slowly at the initial stages, gradually increase in fraction and are most abundant at carrying capacity (Fig 3, blue line in central panel). On the other hand, mutants with larger death rates grow relatively fast at the initial stages, and subsequently decrease in percentage, when the system reaches carrying capacity (Fig 3, yellow line in central panel). In other words, if the mutants are characterized by decreased divisions, we expect to observe the largest fraction of mutants when the patch reaches its maximum population; in contrast, if the mutants are characterized by increased deaths, then the percentage of mutants is larger at intermediate stages of growth compared to patches that reach capacity.

**Fig 3.**
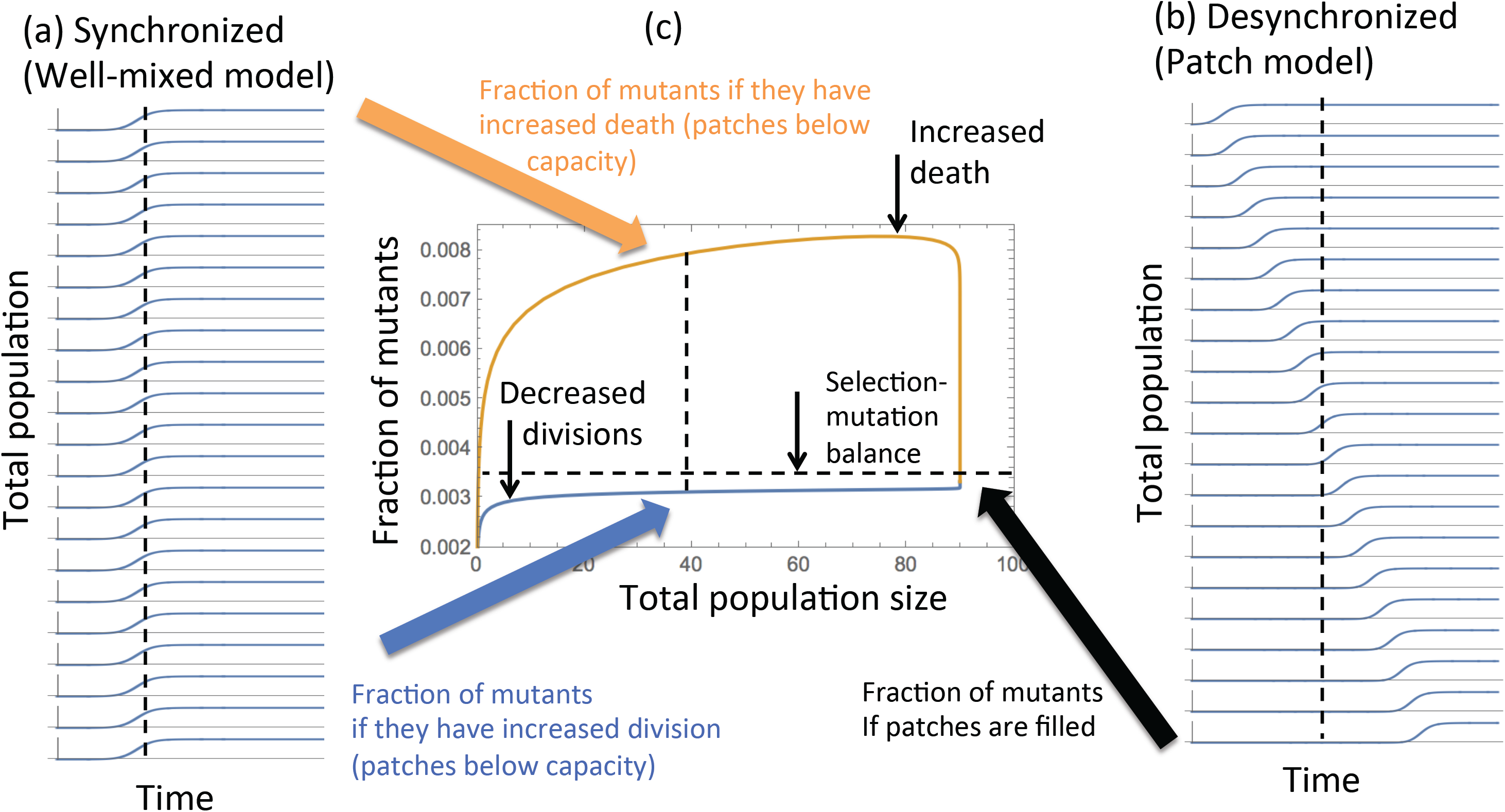
Disadvantageous mutants – an intuitive picture. (a) A schematic representing total population time-series in different patches in a patch model. (b) Same for the well-mixed model represented as a collection of identical, synchronous patches. At the same total population size, in a patch model some populations are at carrying capacity, and some are zero, while in the well-mixed model, all the “patches” are partially filled. (c) The fraction of mutants (characterized by increased death and by decreased divisions) as a function of the population size. Patches with populations below carrying capacity have more mutants than patches at carrying capacity, if the mutants are characterized by increased death. Patches with populations below carrying capacity have fewer mutants than patches at carrying capacity, if the mutants are characterized by decreased divisions.

Next we note that a well-mixed system can be viewed as a superposition of identical, independent smaller patches that all grow simultaneously (figure 3(a)). A (proper) patch model is also a collection of patches, but the growth in different patches does not happen simultaneously; instead, it starts in one patch, after a while a second patch starts growing, etc (Fig 3(b)). Therefore, an important difference between the well-mixed system and a patch system is that in the latter model, different patches are desynchronized, such that at a given point in time some patches are completely filled to capacity while others have not started growing yet.

Keeping this in mind, we can see whether a synchronized (well-mixed) or a desynchronized (patch) model will contain a larger number of mutants. If the mutants have decreased division rates and their percentage grows with total population size, then we are likely to find more mutants in a desynchronized system (spatial or fragmented) that consists of a number of full patches (maximum size, maximum mutant percentage), plus a number of empty patches that do not contribute. In a synchronized (i.e. mixed) system, populations in all patches will lie below carrying capacity at total size M, resulting in fewer mutants. On the other hand, if the mutants have increased death rates and their percentage is larger at the intermediate stages of population growth, then we expect to have more mutants in a fully synchronized system (i.e. a mixed system), which is equivalent to a set of identical patches that are all relatively early in their growth (and thus contain a relatively large percentage of mutants). In the desynchronized (spatial or fragmented) system, populations in several patches will have reached carrying capacity when the total population size reaches M, and thus will have already experienced a decline in mutant percentages.

## Growth laws for neutral, advantageous, and disadvantageous mutants in spatial and non-spatial models

Observations presented so far can be generalized by deriving growth laws of mutants in different scenarios, see Table 1 and Section 4 of the Supplement for details.

**Table 1.** The growth laws of mutants in different spatial and non-spatial growth scenarios, for disadvantageous, neutral, and advantageous scenarios. α is a parameter that quantifies the mutant advantage, α=(L_m_-D_m_)/(L_w_-D_w_).

|  | 1D | 2D | 3D | Exponential |
| --- | --- | --- | --- | --- |
| Disadvantageous | $uN$ | $uN$ | $uN$ | $uN$ |
| Neutral | $uN^2$ | $uN^{3/2}$ | $uN^{4/3}$ | $uN \ln N$ |
| Advantageous | $uN^3$ | $uN^2$ | $uN^2$ | $uN^{(2\alpha-1)/\alpha}$ |

Consider a one-dimensional growth, where the population spreads in one direction (examples of such growth can be found in the geometry of colonic crypts [22,23], or in mitotic zone germ cells in Caenorhabditis elegans [24]). The mathematical abstraction of this process is a cylinder, which is a rectangular domain of width W, with the initial cell configuration aligned along one of the boundaries and periodic boundary conditions imposed in the transversal direction. The cell population in this case will engage in a linear growth (such that the mean total population N=W*L grows as N∼t). The number of disadvantageous mutants in this setting will scale with the total population as specified in the first column of Table 1, as these mutants will typically form finite “bubbles” and thus their number will be entirely driven by production. If mutants are neutral, then on average, each newly created mutant will give rise to a clone that grows linearly in time, thus giving a quadratic growth law (uN^2^), see Figure 4, curves (a,b). Finally, advantageous mutants, when created, will form expanding clones whose width will grow as the colony proceeds to expand; in other words, advantageous colonies comprise (on average) increasing fractions of the total population size, adding an extra power of N to the growth law (uN^3^), see Figure 4, curves (c-g). The number of neutral and advantageous mutants in a 1D colony of a fixed size negatively correlates with the cylinder width: the number of mutants is inversely proportion to the first power of width, W, for neutral, and to the second power of W for advantageous mutants, see Section 4.1 of the Supplement. Note that in the extreme case where W=1, we have a one-dimensional growing array of cells. In this special case [23], in the absence of cell death, all mutants regardless of their fitness properties behave as uN^2^.

**Fig. 4.**
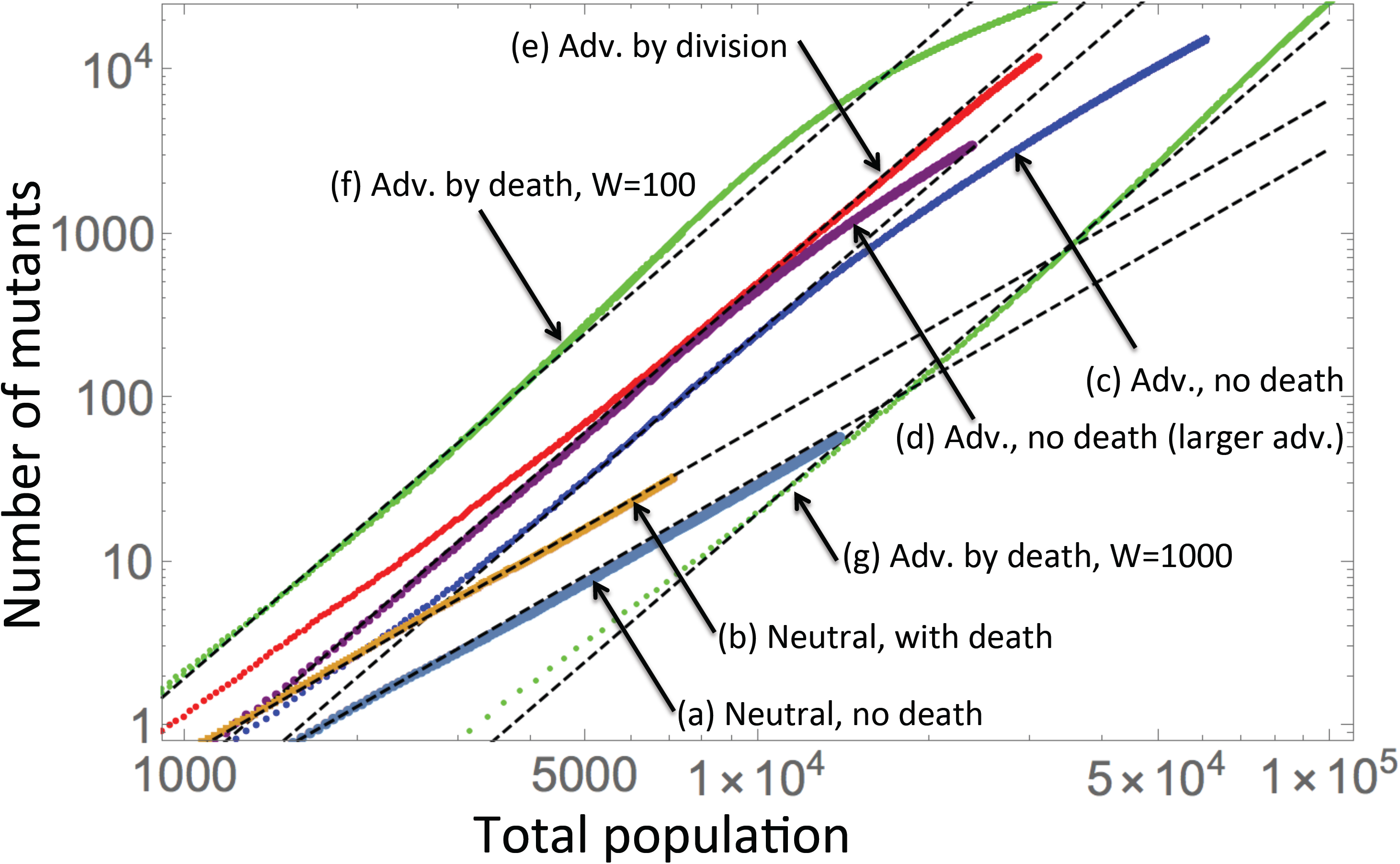
Mutants in the cylindrical geometry: the number of mutants as a function of the total population, averaged over 1000 runs. Cases (a,b) are neutral, and the corresponding dashed black lines have slope 2 in the log-log plot. Cases (c-g) are advantageous, and the dashed lines have slope 3. The following parameters are used: (a) Neutral, no death: L_w_=L_m_=0.7, D_w_=D_m_=0. (b) Neutral, with death: L_w_=L_m_=0.7, D_w_=D_m_=0.2. (c) Advantageous, no death: L_w_=0.7, L_m_=0.9, D_w_=D_m_=0. (d) Advantageous, no death, larger advantage: L_w_=0.7, L_m_=1.0, D_w_=D_m_=0. (e) Advantageous by division, with death: L_w_=0.7, L_m_=0.8, D_w_=D_m_=0.2. (f) Advantageous by death: L_w_=L_m_=0.7, D_w_=0.2, D_m_=0.1. (g) Same as (f), but with a wider cylinder: W=1000. The rest of the parameters are u=5×10^−5^, W=100 (except (g)).

Next, consider range expansion in 2D (e.g. yeast colony expansion [25], 2D melanoma cultures [26,27], where the population grows outward as an expanding circle. In this case, the total population follows the so-called surface growth: N∼t^2^. Mutant cells behave as specified in the second column of Table 1. In particular, disadvantageous mutants are again proportional to the total population; neutral mutations are expected to give rise to colonies whose size does not increase or decrease as a fraction of the total population (the 3/2 law), Figure 5(a,b); advantageous mutations create expanding colonies (the quadratic law), Figure 5(c-f). In a 3D range expansion, which is relevant for example for most solid tumors or biofilms [28], the total population engages in a 3D surface grows as N∼t^3^. Mutants behave as described in the 3^rd^ column of Table 1, and further details are provided in Section 4.2 of the Supplement.

**Fig. 5.**
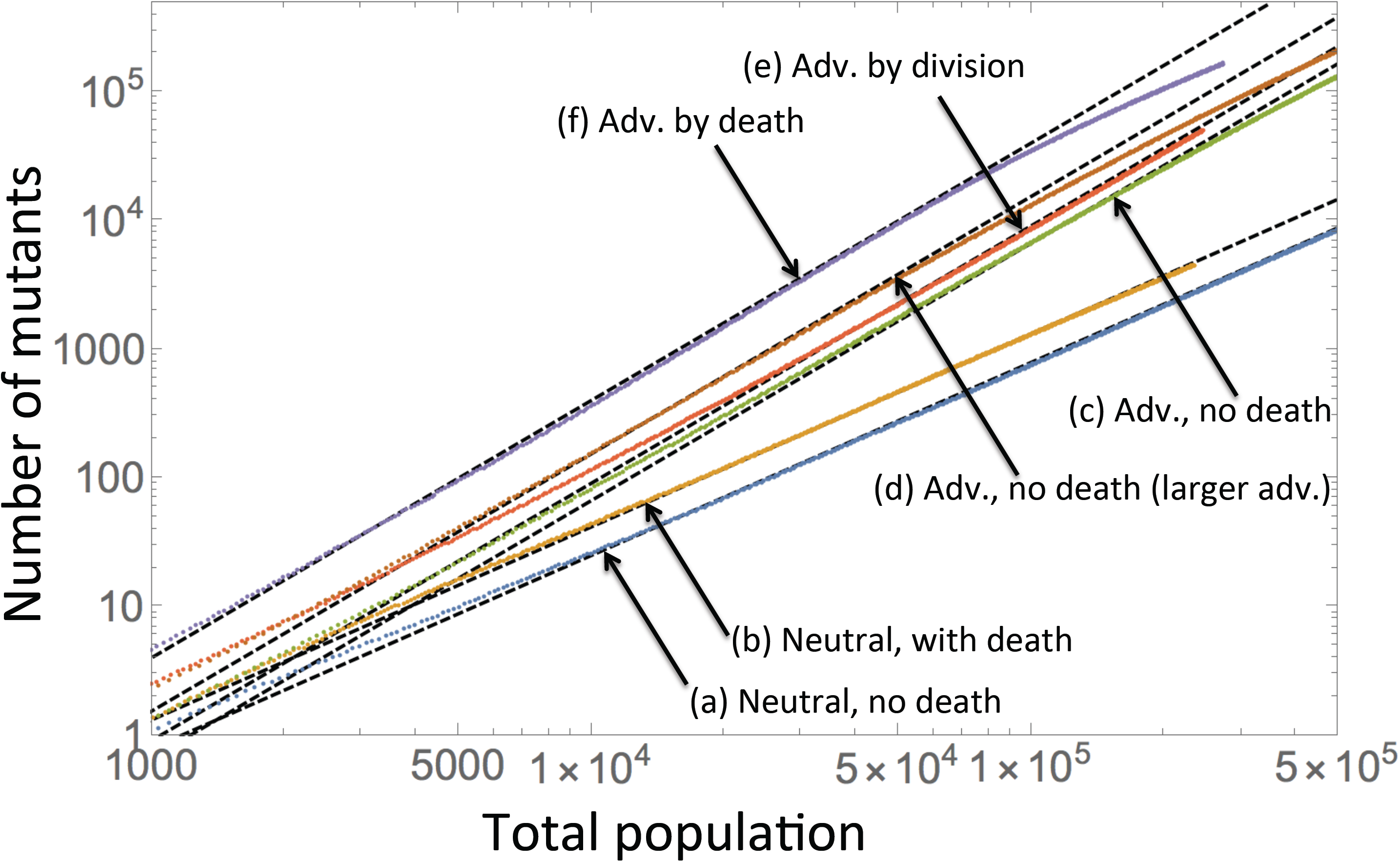
Mutants in the circular geometry: the number of mutants as a function of the total population, averaged over 1000 runs. Cases (a,b) are neutral, and the corresponding dashed black lines have slope 3/2 in the log-log plot. Cases (c-f) are advantageous, and the dashed lines have slope 2. The following parameters are used: (a) Neutral, no death: L_w_=L_m_=0.7, D_w_=D_m_=0, 2000 runs. (b) Neutral, with death: L_w_=L_m_=0.7, D_w_=D_m_=0.2, 1366 runs. (c) Advantageous, no death: L_w_=0.7, L_m_=0.9, D_w_=D_m_=0, 2000 runs. (d) Advantageous, no death, larger advantage: L_w_=0.7, L_m_=1.0, D_w_=D_m_=0, 2000 runs. (e) Advantageous by division, with death: L_w_=0.7, L_m_=0.8, D_w_=D_m_=0.2, 1426 runs. (f) Advantageous by death: L_w_=L_m_=0.7, D_w_=0.2, D_m_=0.1, 1396 runs. The mutation rate is u=5× 10^−5^.

For comparison, results for non-spatial, exponential growth were derived, for example, by [29] and are given in the last column of Table 1. The growth of advantageous mutants in an exponentially expanding population is given by M(exp, adv)∼uN^(2α-1)/α^, where α is a parameter that quantifies the mutant advantage (α=L_m_-D_m_)/(L_w_-D_w_)). Note that as α-> ∞, we have at most M(exp, adv)∼uN^2^, and for all finite values of fitness advantage, the power is less than 2.

These laws are valid under some restrictions specified in Section 4 of the Supplement. In particular, the laws for advantageous mutants hold for small mutant advantage, and also on the time scales before all the cells in a growing colony’s front are replaced by mutants. In the long-term, the replacement of all cells by advantageous mutants is an inevitable outcome in the presence of death, and an approximate outcome as t-> ∞ even in the absence of death. Once this happens, the growth law will be M∼N.

The laws of Table 1 are very general and hold in the presence and in the absence of cell death, and also in the presence and in the absence of back mutations (see Materials and Methods). The proportionality coefficients depend on particularities of the underlying dynamics (for example the type of grid used and the number of neighbors, as well as the death to division ratios), but the power laws are universal.

The growth laws derived here have direct consequences for the expected numbers of mutants in equally sized populations growing in different dimensions (and mass-action). The proportion of neutral mutants scales as uN in 1D, uN^1/2^ in 2D, uN^1/3^ in 3D, while it is u log N in exponentially growing populations (see also [12]). That is, the number of neutral mutants is always larger in spatial systems compared to the well-mixed system. In space, the proportion of neutral mutants is the largest in low dimensions.

For advantageous mutants, the proportion of mutants is given by uN^2^ in 1D, uN in 2D, and uN in 3D, while it is uN^(α-1)/α^ in exponentially growing populations. Again, it decreases with the dimensionality of the system, and is always the smallest in mass-action.

Finally, for disadvantageous mutants, the power law of mutant growth is the same in all dimensions (and is given by uN). Therefore the results are more subtle and depend on the particular setup. As was shown in the previous section, the behavior depends on whether the disadvantage is manifested through differences in division or death rates.

## Discussion and Conclusion

We have used computational models to study mutant evolution in spatially structured and fragmented populations, focusing on the average number of mutants present when the total population size has reached a threshold. Previous work [12] established that for neutral mutants, spatial restriction results in a larger number of mutants that are present in a population of a defined size. This was attributed to jackpot events occurring even at relatively large populations sizes in spatially structured populations due to the occurrence of range expansion. In contrast, jackpot mutation events can only occur at very early stages of growth in mixed systems. We extended this analysis by additionally considering advantageous and deleterious mutants, in the absence and presence of cell death. While the results for advantageous mutants were similar to those for neutral mutants (more mutants in spatial than mixed systems), we found the opposite trend for disadvantageous mutants. If the extent of the disadvantage was sufficiently large, the number of mutants in spatially structured populations was *smaller* than in mixed systems. This has important practical implications, for example for understanding the presence of drug resistant mutants prior to the start of treatment in cancers [30] or bacterial populations that form a biofilm [31]. Spatial structure is predicted to make it less likely that mutants are present before treatment is started, and if they are present, their average numbers are lower in spatially structured compared to mixed systems. Drug resistant mutants have often been shown to be characterized by a fitness cost compared to drug-sensitive cells [32,33], and a better understanding of spatial mutant evolution is important for our ability to understand and predict treatment outcome. More generally, we derived growth laws for different types of mutants in different settings that vary in the extent of the spatial restriction experienced by cells, and these laws may aid in predicting mutant burdens in growing spatial populations.

Another interesting result was our finding that qualitatively identical results are obtained if we consider evolutionary dynamics in fragmented rather than spatially structured populations. The same outcomes were obtained for a patch model where individuals in each patch could migrate to any randomly chosen patch in the system (and not just the neighboring patches). Therefore, the properties of mutant evolution described here and also in [12] might not be a particular property of spatial systems, but more generally of fragmented systems. Our intuitive explanation for the differences in the numbers of disadvantageous mutants does not rely on any spatial restrictions in cell migration, but rather on the de-synchronization of mutant dynamics in the patches of a fragmented system. Unlike a well-mixed system, jackpot events in a fragmented system can occur relatively late in the process and still make a significant difference (regardless of the patches’ spatial arrangement or the absence thereof). An analogous result has been reported for predator-prey type dynamics: the properties of a 2-dimensional agent based model were similar to those in a fragmented patch model, and a similar connection was made between a spatially structured and a fragmented habitat [34].

Overall this study has demonstrated complex evolutionary dynamics in populations that are not well-mixed. We demonstrated that evolution can be influenced in different ways by spatial structure or habitat fragmentation, depending on the relative fitness of the mutant and depending on the parameter through which the fitness difference is expressed. These results can also guide future experiments to address some of the computational observations reported here. Experimental results from 2-dimensional spatial growth of cells, such as reported in [12], should be compared to analogous results from a fragmented system, for example where cells are grown in a collection of different wells and periodically transferred to other, randomly chosen wells. This could test our prediction that the two scenarios follow the same evolutionary laws. On a more complex level, it would be interesting to devise an experimental system where the evolutionary dynamics of disadvantageous mutants is studied, comparing scenarios where the disadvantage is brought about by a difference in cell death versus cell division.

## Materials and methods

### Two-dimensional agent-based model

We used a 2-dimensional, agent-based model, where a 2-dimensional square grid is considered. A spot on the grid can be empty or can contain a cell, which is either wild-type or mutant. At each time step, the grid is randomly sampled N times, where N is the total number of cells currently in the system. If the sampled cell is wild-type, the cell attempts division (described below) with a probability L_w_ or dies with a probability D_w_. When reproduction is attempted, a target spot is chosen randomly from the nearest neighbors of the cell (8 neighbors, i.e. the Moore neighborhood, was used unless otherwise noted). If that spot is empty, the offspring cell is placed there. If it is already filled, the division event is aborted. The offspring cell is assigned wild type with probability 1-*u* and it is a mutant with probability *u*. If the sampled spot contains a mutant cell, the same processes occur. Attempted division occurs with a probability L_m_, and the cell dies with a probability D_m_. The offspring of a mutant cell is always a mutant in the absence of back mutation. In a different version of the model, a mutant’s offspring can be of wild-type with probability *u*. Initial and boundary conditions are determined by the specific geometric setting investigated. For 2D spatial simulations, an *nxn* square domain is considered. At the boundaries of the domain, a spot is assumed to have fewer neighbors, i.e more division events will fail. The simulations start with a single wild-type cell, placed into the center of the grid. Simulations always stop before the boundary of the grid is reached. For 1D cylinder simulations, we use an *nxw* rectangular domain of width *w*. We start with an array of *w* wild type cells at the left boundary of the domain, and impose periodic boundary conditions in the transversal direction. In each simulation, the cell population is allowed to grow to a size M, and the number of mutant cells at this size is recorded. Such simulations are performed repeatedly, and the average number of mutants is calculated. Simulation runs, in which the total cell population goes extinct due to stochastic effects, are ignored.

Analysis of 2D spatial stochastic models is presented in Section 3 of the Supplement. Growth laws for different geometries are derived in Section 4 of the Supplement.

### Modeling exponential growth

To compare the spatial dynamics to a well-mixed system, we considered a simple stochastic simulation of exponential growth. Denoting the number of wild-type cells with x_w_ and the number of mutant cells with x_m_, one of the cell types is chosen with a probability given by their proportion in the whole cell population. Wild-types can divide with a probability L_w_ and can die with a probability D_w_. Mutants can divide with a probability l_m_ and die with a probability d_m_. Mutations of wild type cells happen with probability *u*. As in the spatial system, the average number of mutants at population size M was determined.

### A patch model

We also considered an alternative modeling approach to capture mutant dynamics in spatially structured populations. Instead of tracking the spatial location of individual cells, we analyze a model that consists of a two-dimensional grid of nxn patches or demes. Within each patch, local dynamics occur where cells are assumed to mix perfectly. At each time step, cells are allowed to migrate to a different patch with a given rate. In each local patch, Gillespie simulations [35] of the following ordinary differential equation model were run:

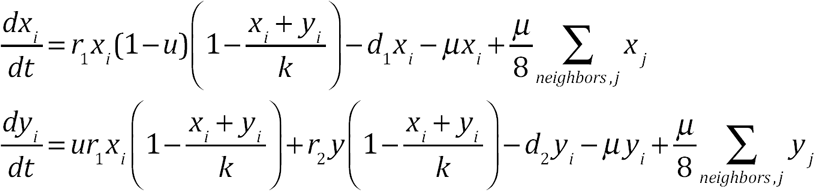

Wild-type cells are denoted by x_i_, and mutant cells by y_i_, where the subscript *i* enumerates the spatial locations in a two-dimensional array. Wild type cells divide with a density-dependent rate r_1_(1-(x+y)/k), die with a rate d_1_, and migrate out of the patch with a rate µ. Migration is assumed to occur to one of the eight neighboring patches, chosen randomly. During replication of the wild-type cells, a mutant cell can be generated with a probability u. Mutant cells divide with a density-dependent rate r_2_(1-(x+y)/k), die with a rate d_2_, and migrate with a rate µ.

In an alternative (fragmentation) model, instead of migrating with rate µ/8 per patch to one of the eight neighboring patches, cells migrate with probability µ/(n-1) per patch to any other patch regardless of its location. This holds for cells in all the patches in the system, thus removing a spatial component from the migration process. Otherwise, the equations are identical to the ones above.

Simulations were started with a single wild-type cell in the middle patch. The simulations were run until the total cell population size, summed over all patches, reached size M. At this point, the number of mutants summed over all patches was recorded. This was done repeatedly, and the average number of mutants at size M was determined. Instances of the simulation that resulted in population extinction across all patches were ignored.

For comparison, Gillespie simulations were performed in a non-fragmented, well mixed system described by the following equations:

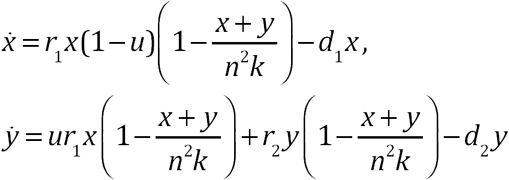

The carrying capacity of the non-fragmented system is taken n^2^ times the carrying capacity of the individual patches (n^2^ is the total number of patches). The average number of mutants at population size M was determined in the same way as in the patch model.

Deterministic (ODE) versions of these models are presented in Section 2 of the Supplement.

