## Supplementary Materials for "Mutant evolution in spatially structured and fragmented populations"

### Mutant dynamics in spatially structured, growing populations

### 1 Simulations: additional results

#### 1.1 Agent-based model simulations

In the main text we report on the abundances of advantageous, neutral, and disadvantageous mutants in spatial and well-mixed system. Some further results are presented in figure S1. In particular, panels (a) and (b) study advantageous mutants. In (a), the advantage is manifested through an increased division rate, and in (b) through a decreased death rate of mutants. Varied is the rate that is unaffected by the mutation under consideration, which is the death rate in (a) and the reproduction rate in (b). As expected, we observe that the number of advantageous mutants is higher in a spatial system (red) compared to the well-mixed system (black) for both types of mutants. The difference increases with death rate (see panel (a)) and decreases with the reproduction rate (panel (b)); in other words, the difference between well-mixed and spatial models is larger for cells with an overall slower expansion rate.

Next, we turn to disadvantageous mutants. In figure S1(c) we study mutants characterized by a lower reproduction rate, and in figure S1(d) the mutants' death rate is higher compared to that for wild type cells. Again, in both panels, the rate unaffected by mutations is varied, and the mutant abundance in spatial (red) and well-mixed (black) system compared at the same population size. As reported in the main text, we observe that mutants with a lower reproduction rate are more abundant in a spatial model. As seen in panel (c), the differences becomes smaller with an increased death rate of cells.

Finally, we turn to mutants characterized by a larger death rates. It is reported in the main text that if the disadvantage is sufficiently pronounced, we expect to find more such mutants in a well-mixed system compared to a spatial system. This is what we see in figure S1(d), when the reproduction rates are lower than a threshold. This trend reverses, however, when the reproduction rates become higher. This provides further information about the phenomenon reported in the main text. The mutant disadvantage must be sufficiently high, for the well-mixed system to accumulate more mutants than the spatial system, and this advantage is measured against the background reproduction rate of the cells. As the reproduction rate gets higher, the difference between mutant and wild type death rates must also become higher, to observe more mutants in a well-mixed system compared to the spatial system.

#### 1.2 Patch model simulations

We have run numerical simulations of the stochastic patch model to determine whether spatial arrangement of patches makes a difference for the abundance of mutants at a given system size. Figure S2 shows the results for neutral (a) and advantageous (b) mutants, while figure 2 of the main text contains information about disadvantageous mutants. In figure S2, two types of the patch model are compared. In the spatial 2D patch model (dark green bars), we used a 2D array of patches, where cells could migrate only between neighboring

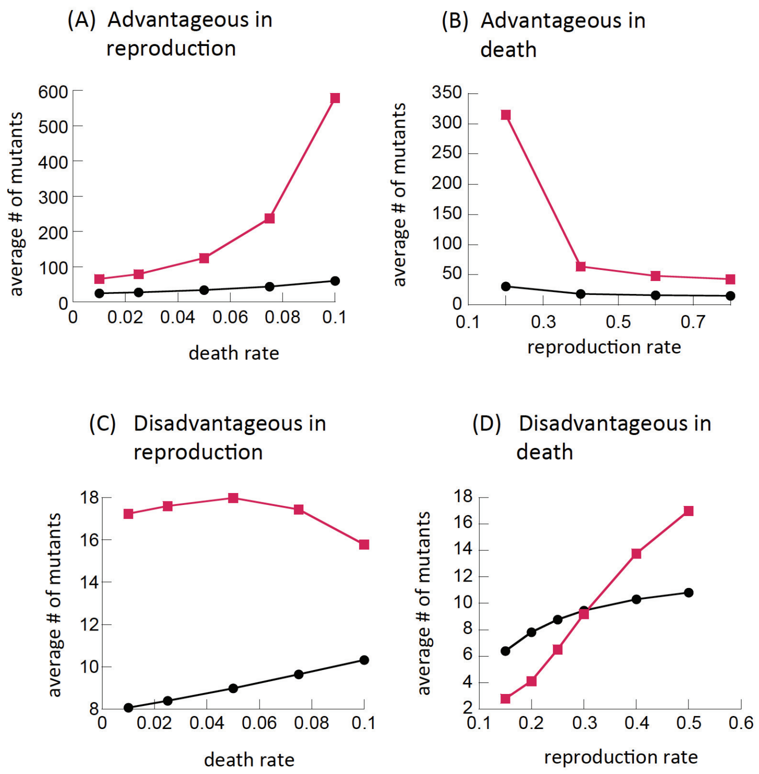

Figure S1: The abundance of mutants in spatial 2D simulations (red) and in a well-mixed system (black), as a function of parameters. (a) Mutants have a larger reproduction rate ($L_w = 0.2, L_m = 0.25$); varied is the death rate (equal for mutants and wild types). (b) Mutants have a smaller death rate ($D_w = 0.1, D_m = 0.09$); varied is the reproduction rate (equal for mutants and wild types). (c) Mutants have a smaller reproduction rate ($L_w = 0.25, L_m = 0.2$); varied is the death rate (equal for mutants and wild types). (d) Mutants have a larger death rate ($D_w = 0.05, D_m = 0.1$); varied is the reproduction rate (equal for mutants and wild types). The rest of the parameters are as in figure 1 of the main text.

patches (with each patch having 8 neighbors as in the Moore model). In the "fragmentation model" (light green bars) migration could happen between any patches regardless of their position. In the latter case, the model cannot be regarded as spatial per se. Nonetheless, similar trends were observed in both models. For advantageous and neutral mutations, more mutants were observed in the patch model (a 2D or a fragmented, non-spatial patch system) compared to the well-mixed system with the same total number of cells; the effect is less pronounced but still clearly present in the fragmented system.

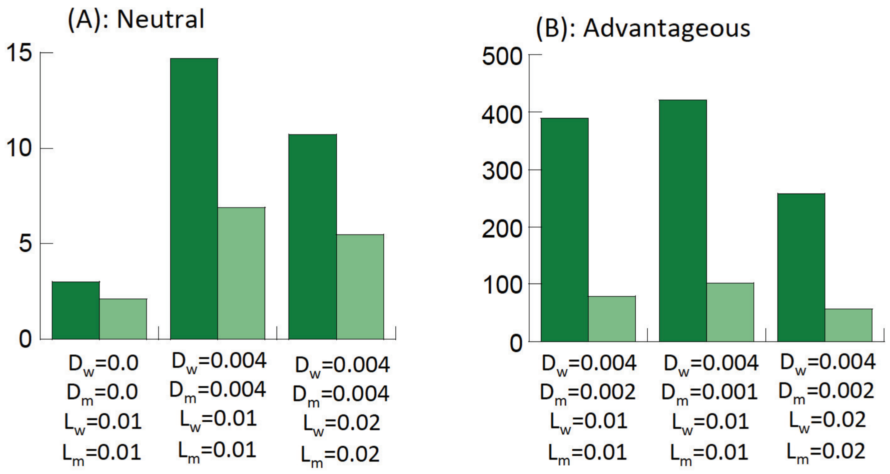

Figure S2: Comparison of the number of mutants in stochastic patch model simulations and a well-mixed system. The bars represent the ratio between the mean number of mutants in the patch model and the mean number of mutants in well-mixed systems at equal size, $N = 10^4$. Dark green bars correspond to the 2D patch model with 8 nearest neighbor migration. Light green bars correspond to the fragmented model where migration happens to all patches. (a) Neutral mutants. (b) Advantageous mutants. Between $10^6$ and $4 \times 10^7$ runs were performed for each bar. Division and death rates are indicated below each bar. Other parameters are $u = 2 \times 10^{-5}$, $k = 100$, $\mu = 10^{-5}$, $n = 31 \times 31$ patches.

### 2 Disadvantageous mutants: a deterministic metapopulation model

#### 2.1 Basic formulations and selection mutation balance

Let us denote the wild type population as $x(t)$ and the mutant population as $y(t)$. Denote the rate of mutations as $u$, the division and death rates of wild type cells as $L_x$ and $D_x$, and the division and death rates of mutants as $L_y$ and $D_y$. Then the competition dynamics of cells can be formulated as follows:

$$\dot{x} = L_x x(1-u)\left(1 - \frac{x+y}{\mathcal{K}}\right) - D_x x, \quad (1)$$

$$\dot{y} = (L_x x u + L_y y)\left(1 - \frac{x+y}{\mathcal{K}}\right) - D_y y, \quad (2)$$

where $\mathcal{K}$ is the carrying capacity. If the mutants are disadvantageous, that is, if

$$\nu = \frac{D_y}{D_x} - \frac{L_y}{L_x} > 0, \quad (3)$$

then the selection mutation balance predicts that the equilibrium number of wild type cells is given by

$$\bar{x} = \mathcal{K}\left(1 - \frac{D_x}{L_x}\right), \tag{4}$$

and the number mutants is

$$\bar{y} = \bar{x}u\left(\frac{D_y}{D_x} - \frac{L_y}{L_x}\right)^{-1} \tag{5}$$

(we only took the largest contributions in terms of small $u$).

In model (1-2), as the population approaches the carrying capacity, the divisions slow down, while deaths remain occurring at a constant rate. We will refer to this model as *division-controlled growth*, which follows the terminology of [1]. Alternatively, we can assume that as the population grows, the death rate increases, while the division rate stays constant. In the mass action case, this can be modeled as follows:

$$\dot{x} = L_x x(1-u) - D_x x\left(1 + \frac{x+y}{\mathcal{K}}\right), \tag{6}$$

$$\dot{y} = (L_x xu + L_y y) - D_y y\left(1 + \frac{x+y}{\mathcal{K}}\right). \tag{7}$$

We will refer to this model as *death-controlled growth* [1]. In this case, the equilibrium population size is

$$\bar{x} = \mathcal{K}\left(\frac{L_x}{D_x} - 1\right), \tag{8}$$

and the number of mutants in selection mutation balance is again given by formula (5).

The early dynamics of wild type and mutant populations in models (1-2) and (6-7) are identical. Assuming that $x + y \ll \mathcal{K}$, we can solve the resulting linear equations exactly, to obtain

$$x_{lin}(t) = e^{(L_x - D_x)y}, \quad y_{lin}(t) = L_x u \frac{e^{(L_x-D_x)t} - e^{(L_y-D_y)t}}{(L_x - D_x) - (L_y - D_y)}. \tag{9}$$

#### 2.2 Decreased divisions and increased death

A mutant is disadvantageous if inequality (3) holds. If we fix $L_x$ and $D_x$, the division and the death rates of the wild type cells, this inequality defines a half plane in the $(L_y, D_y)$ space, where mutants are disadvantageous (more precisely, it is the region above the line $D_y/L_y = D_x/L_x$, see figure S6, the red line). Note that this definition is not equivalent to using the linear growth rate to define fitness, because instead of linear initial expansion of the mutants, it measures their steady state level in the presence of the wild types. For comparison, the line where the linear growth rates of mutant and wild-type cells are equal to each other, is shown in figure S6 in green.

The quantity $\nu$ defined in (3) can be seen as a measure of fitness disadvantage. Mutants with equal $\nu$ have the same level of disadvantage, which is for the purposes of this paper defined as the same level of the selection mutation balance equilibrium. The definition of $\nu$ (equation (3)) with fixed $L_x, D_x$ generates a one-parametric family of types of equal disadvantage. Disadvantage can be achieved by different combinations of division and death rates, $L_y, D_y$. In particular, if the mutants have the same death rates as the wild types, and their disadvantage is achieved through lowered division rates, then we have a type with **decreased divisions**,

$$L_y = L_x(1-\nu), \quad D_y = D_x; \tag{10}$$

in figure S6, "decreased division" mutants correspond to all the points on the horizontal dashed line (in the disadvantageous region). If on the other hand, the division rate of mutants matches that of the wild types, we have a type with **increased death**:

$$L_y = L_x, \quad D_y = D_x(1+\nu); \tag{11}$$

in figure S6, "decreased death" types correspond to disadvantageous points on the vertical dashed line. Note that if the two types have the same fitness (i.e. converge to the same selection-mutation balance), the percentage decrease in the division rate must be equal to the percentage increase in the death rate. The linear growth rate of the two types is however different, and is given by

$$L_y - D_y = \begin{cases} L_x - D_x - \nu L_x, & \text{decreased divisions,} \\ L_x - D_x - \nu D_x, & \text{increased death.} \end{cases}$$

We can see that since $L_x > D_x$, the "increased death" mutants are always characterized by a faster growth compared to the "decreased divisions" mutants.

Figure S3(a) shows an example of the numbers of mutants under the assumptions of decreased divisions (blue) and increased deaths (yellow); specific parameter values are given in the figure caption. The behavior of the wild types can be seen from the dashed green line that shows $x(t)/1000$. Before the population reaches the carrying capacity, the "increased death" mutants grow faster than the "decreased divisions" mutants. At later times, they both reach the same selection mutation equilibrium.

A useful representation of this information is given in panel (b) of figure S3, where we plot the number of mutants, $\mathcal{M}(x)$, contained in the system of size $x$. This quantity is presented for both types (decreased divisions and increased death) in figure S3(b).

#### 2.3 The patch model

Consider $N$ patches, such that in each patch cells undergo deterministic mass-action dynamics of divisions and deaths, subject to a carrying capacity $K$. We

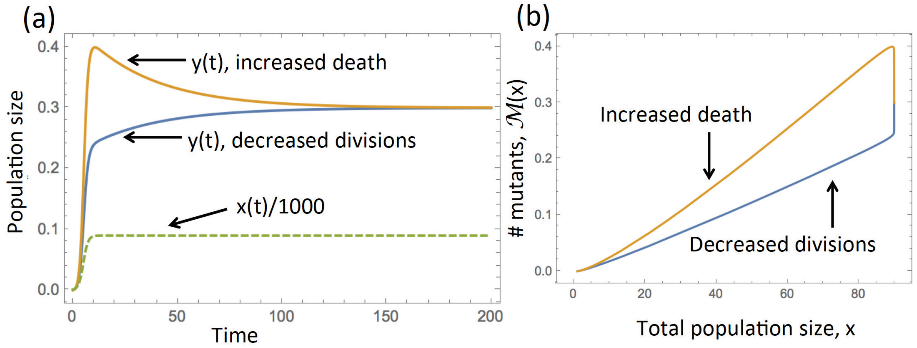

Figure S3: Solutions of system (1-2), where the division and death rates of the wild types are fixed, and the mutant rates are given by equation (10) for the "decreased divisions" type, and equation (11) for the 'increased death" type (only one type of mutant is included at a time). (a) The level of mutants as a function of time for the two systems; the wild types are given by the green dashed line, where the value was divided by 1000 to bring it to the same scale. (b) The number of mutants plotted as a function of the total cell population for the two types. The parameters are $L_x = 1; D_x = 0.1, \nu = 0.3, u = 0.001, K = 100$. The fitness disadvantage is $\nu = 0.3$.

assume that patches communicate with each other through migrations. Let us denote the migration matrix as $M = \{M_{ij}\}$, where $M_{ij}$ is the probability that, given that a cell from patch $i$ migrates, it is transferred to patch $j$; we have $\sum_{j\neq i} M_{ij} = 1$ for all $i$. We explore two types of such matrices:

1. **A 1D spatial model:** a ring of patches where only migration between nearest neighbors is possible:

$$M_{ij} = \begin{cases} 1/2, & |i-j| = 1, \\ 0, & \text{otherwise}, \end{cases} \tag{12}$$

   with the additional nonzero values $M_{1N} = M_{N1} = 1/2$.

2. **A complete graph model**: all patches are connected:

$$M_{ij} = \begin{cases} 1/(N-1), & i \neq j, \\ 0, & i = j. \end{cases}$$

The ODEs in each patch are given by equations similar to (1-2) for division-controlled growth, or (6-7) for death-controlled growth, with $\mathcal{K} = K$ and migration terms included. In the former case, the system at each patch looks like this:

$$\dot{x}_i \quad = \quad L_x x_i (1-u)\left(1 - \frac{x_i + y_i}{K}\right) - D_x x_i - \mu\left(x_i - \sum_{j=1}^{N} M_{ji} x_j\right), \tag{13}$$

$$\dot{y}_i = (L_x x_i u + L_y y_i)\left(1 - \frac{x_i + y_i}{K}\right) - D_y y_i - \mu\left(y_i - \sum_{j=1}^{N} M_{ji} y_j\right), \quad (14)$$

where $x_i$ and $y_i$ are the numbers of wild type and mutant cells in patch $i$, respectively. The initial value problem is completed with the following initial condition:

$$x_i(0) = \begin{cases} 1, & i = (N+1)/2, \\ 0, & \text{otherwise}, \end{cases} \qquad y_i(0) = 0 \quad \forall i \quad (15)$$

(we assumed an odd number of patches), that is, there is a single wild type cell in the middle patch, and the rest of the patches are unoccupied.

The results of the $N$-patch model will be compared with an unfragmented, mass-action system of size (carrying capacity) $\mathcal{K} = NK$:

$$\dot{X} = L_x X(1-u)\left(1 - \frac{X+Y}{KN}\right) - D_x X, \quad (16)$$

$$\dot{Y} = (L_x X u + L_y Y)\left(1 - \frac{X+Y}{KN}\right) - D_y Y, \quad (17)$$

$$X(0) = 1, \quad Y(0) = 0. \quad (18)$$

For death-controlled growth, all equations are modified accordingly.

#### 2.4 The number of mutants in fragmented and mass-action systems

Let us compare the number of mutants obtained in the fragmented system with migrations and in the mass action system of the same total carrying capacity. Suppose we are interested in measuring the number of mutant at a total population size $N_{tot}$. Then we define the time $t_{fr}$ and $t_{ma}$ (for "fragmented" and "mass-action" respectively) as follows:

$$\sum_{i=1}^{N} (x_i(t_{fr}) + y_i(t_{fr})) = N_{tot}, \quad X(t_{ma}) + Y(t_{ma}) = N_{tot}.$$

We want to compare the numbers of mutants,

$$M_{fr}(N_{tot}) = \sum_{i=1}^{N} y_i(t_{fr}), \quad M_{ma}(N_{tot}) = Y(t_{ma}).$$

Figures S4(a) and S5(a) show two examples of functions $M_{fr}(N_{tot})$ and $M_{ma}(N_{tot})$, for "decreased divisions" and "increased death" mutants respectively. We obtained numerical solutions of systems (13-14) with initial conditions (15) and the migration matrix (12) (a 1D ring of patches), to plot $M_{fr}$ (the blue line) as a function of the total population size. The corresponding number of mutants

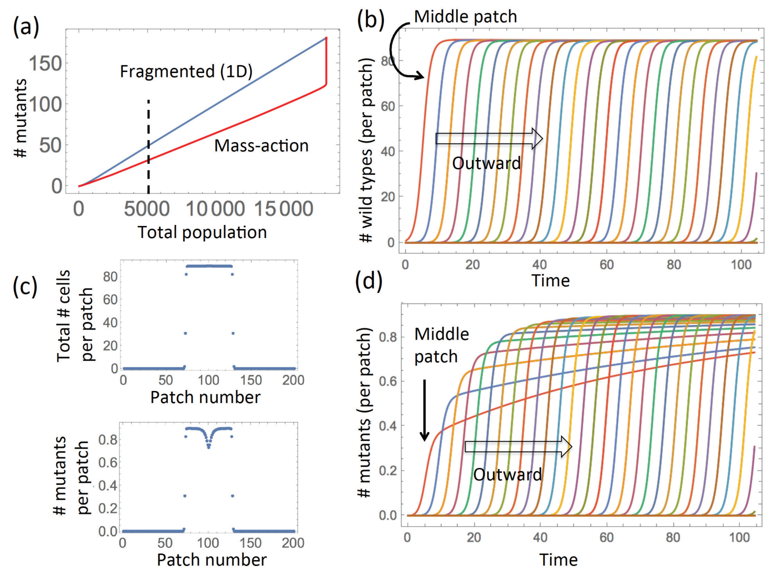

Figure S4: Mutant dynamics in the case of "decreased divisions" mutants. (a) Comparison of the mutant number as a function of the population size for the fragmented (blue) and mass-action (red) systems. A particular value of the total population size, $N_{tot} = 5000$, is indicated by a dashed vertical line. (b) The wild type populations in different patches as functions of time, for $t \in [0, t_{fr}]$ corresponding to $N_{tot} = 5000$. The leftmost line corresponds to the middle patch, and the rest of the lines to consecutive patches moving outward. (d) Same as (b) for the mutant populations. (c) The total populations (top) and the mutant populations (bottom) in all the patches at time $t_{fr}$. The parameters are $N = 201$ patches, $L_x = 1; D_x = 0.1, L_y = 0.9; D_y = 0.1, u = 0.001, K = 100; \mu = 0.01$.

in the mass action system, $M_{ma}$ (the red line), was obtained from system (16-18). We observe that in figures S4(a) ("decreased divisions"), there are more mutants in the fragmented, spatial system, and in figures S5(a) ("increased deaths"), there are more mutants in the mass action system.

Panels (b,c,d) of figures S4 and S5 elucidate some underlying patterns. Panels (b,d) depict population dynamics in the patches. The initial wild type population in the "middle" patch number 101 (out of 200) grows and "seeds" the neighboring patches, whose population starts increasing, which in turn gives rise to growth in the next patches, etc, see figures S4(b) and S5(b). Because of the symmetries of the ODEs, patches equidistant from the middle have identical solutions. Figures S4(d) and S5(d) show the numbers of mutants as they evolve in time, in each patch. Again, the leftmost line corresponds to the middle patch that was occupied initially. Individual trajectories of mutants in patches behave differently: "decreased divisions" mutants (figure S4(d)) grow monotonically toward the mutation selection balance (compare the blue line in figure S3(a)); "increased death" mutants (figure S5(d)) "overshoot" and then decrease toward the mutation selection balance (compare the yellow line in figure S3(a)).

Figure S6 generalizes these results. It presents the parameter space with coordinates $L_y$ and $D_y$, where the points where $M_{ma}(N_{tot}) = M_{fr}(N_{tot})$ are shown as a purple line (parameters other than $L_y$ and $D_y$ are fixed). In the region above the purple line (gray shadowing), there are more mutants in the mass action system, and below it there are more mutants in the fragmented system. In particular, the system of figure S4 has coordinates $(L_y, D_y) = (0.9, 0.1)$ and falls outside the gray region, and the system of figure S5 corresponds to $(1, 0.2)$ and is inside the gray region; both points are marked by a "$*$" in the figure. In fact, we can see that all the "decreased divisions" mutants (points that lie on the dashed horizontal line in the disadvantageous region) fall under the purple line, that is, decreasing division rate leads to having more mutants in the fragmented system. It is somewhat less straightforward with "increased death" mutants (disadvantageous mutants on the dashed vertical line): if the increase in death is sufficiently large, then such points are above the purple line (and there are more mutants in mass action). For a small increase in death rates, there are more mutants in the fragmented system.

#### 2.5 A simplified theory of mutant abundance in spatial and mass-action systems

Simulations in panels (b,d) of figures S4 and S5 were run for time $t_{fr}$ that corresponds to the fragmented system reaching a specific total size $N_{tot} = 5000$, denoted by the dashed vertical line in panel (a). At that time, the total numbers of cells per patch and the numbers of mutants per patch are shown in panel (c). We observe that at time $t_{fr}$,

(i) not all patches are occupied,

(ii) in most occupied patches, except for the outmost ones, the total populations have reached the carrying capacity (given by $L_x(1 - D_x/L_x)$), and

(iii) in most patches, the number of mutants is at the selection mutation balance. The exceptions are again the outmost patches that have not stabilized yet, and the middle patches, where the growth of mutants is slower.

Using the observations of patch dynamics listed above, let us approximate the number of mutants in a 1D fragmented system. At the time the total population reaches a size $N_{tot}$, we assume that there will be a number of patches that are completely occupied, with population at carrying capacity, $\bar{x} = K(1 - D_x/L_x)$, and a number of patches that have not been reached yet. There are

$$n = \frac{N_{tot}}{K(1 - D_x/L_x)}$$

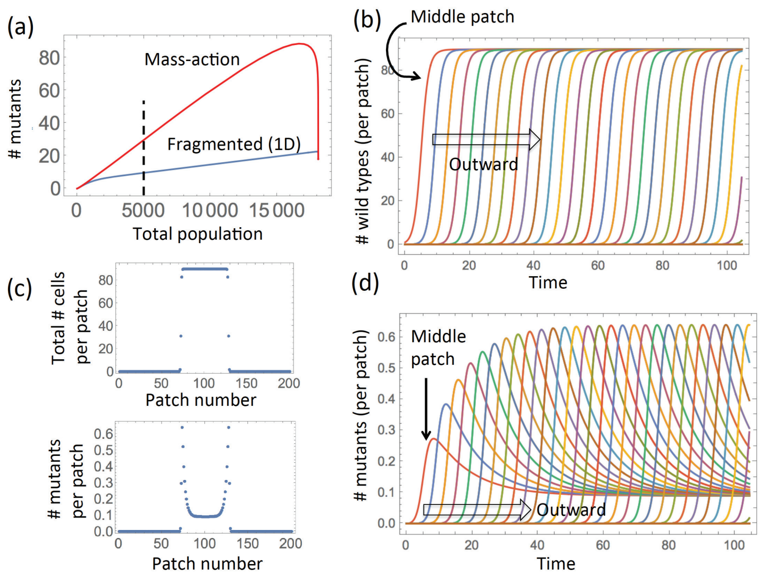

Figure S5: Mutant dynamics in the case of "increased death" mutants. Same as in figure S4, except $L_y = 1, D_y = 0.2$.

such occupied patches. Therefore, the total number of mutants at size $N_{tot}$ is given by $n\bar{y}$, where $y$ is given by equation (5), and we have

$$M_{fr}(N_{tot}) \approx N_{tot} u \left( \frac{D_y}{D_x} - \frac{L_y}{L_x} \right)^{-1}. \tag{19}$$

To calculate the number of mutants in the non-fragmented population, we use formula (9). We have, using $x_{lin} = e^{(L_x - D_x)t} \approx N_{tot}$, that

$$M_{ma}(N_{tot}) \approx L_x u \frac{N_{tot} - N_{tot}^{\frac{L_y - D_y}{L_x - D_x}}}{(L_x - D_x) - (L_y - D_y)}. \tag{20}$$

The solution set of the equation $M_{fr} = M_{ma}$ using approximations (19, 20) is shown in figure S6 as a black solid line. Although it is similar to the full solution (purple line) for small values of $L_y$, it deviates from it as $L_y$ becomes closer to $L_x$.

One can further simplify formula (20), assuming that $N_{tot} \gg N_{tot}^{\frac{L_y - D_y}{L_x - D_x}}$. Then, solving $M_{ma} > M_{fr}$, one obtains a condition that does not depend of $N_{tot}$:

$$\frac{D_y}{D_x} - \frac{L_y}{L_x} > \frac{(L_x - D_x) - (L_y - D_y)}{L_x},$$

which can be solved for $D_y$ to yield:

$$D_y > D_x. \tag{21}$$

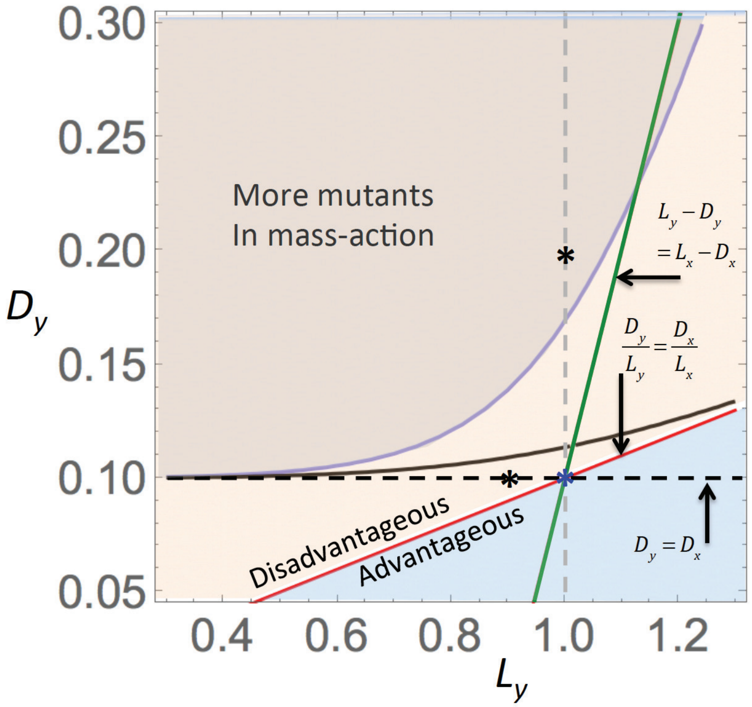

Figure S6: Regions in the parameter space $(L_y, D_y)$, where $M_{ma} > M_{fr}$ (under fixed $(L_x, D_x)$). The red line indicate sets where mutants and the wild types have equal fitness (equation (3) with $\nu = 0$; mutants are disadvantageous above this line); the green line corresponds to equal linear growth rates. The set (21) is above the dashed horizontal line. The solid black line comes from a numerical solution of $M_{ma} = M_{fr}$ with expressions (19,20). The blue solid line is the solution of the same equation using the values obtained from solving the ODEs. $N_{tot} = 1000$, the rest of the parameters are as in figure S4. The point where $L_y = L_x$ and $D_y = D_x$ is marked by a blue star.

In other words, if $D_y > D_x$, there are more mutants in the mass action system, and otherwise there are more mutants in the fragmented system (given that the total sizes of the two populations are the same). Figure S6 shows this approximation as a horizontal dashed line.

#### 2.6 An intuitive explanation

Here we provide an intuitive explanation for the following result:

- If the mutant disadvantage is due to reduced division rates compared to the wild type, then there tends to me more mutants in the fragmented system compared to the mass action system of the same size.

- If the mutant disadvantage is due to increased division rates, then the result is the opposite, and there tends to be more mutants in the mass-action system. This is true only if the disadvantage is sufficiently strong.

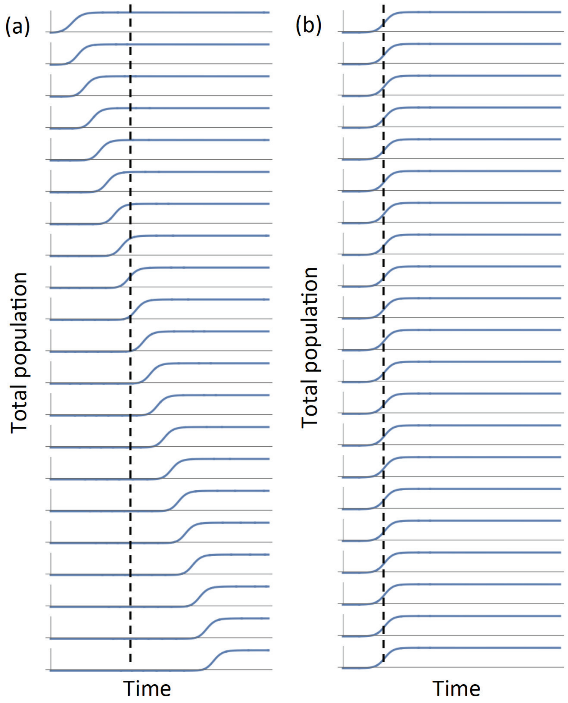

Figure S7: Time series of total populations sizes in patches for (a) a fragmented system with migrations according to a 1D spatial pattern and (b) a number of identical, uncoupled, patches that mathematically are equivalent to the mass action system.

Let us compare two systems: (i) a fragmented system of $N$ patches of carrying capacity $K$ each, where initially there is a single cell in one patch, and populations spread from patch to patch by means of migration, reaching their carrying capacity at different times, and (ii) a mass action system, which can be mathematically represented an a equivalent system of $N$ decoupled identical systems of carrying capacity $K$, with identical initial conditions $x_i(0) = 1/N, y_i(0) = 0$. Figure S7 shows the simulated total population size in a number of patches, for (a) a 1D chain of fragmented patches and (b) a mass-action system represented as $N$ identical patches.

If the target population, $N_{tot}$, is the maximum size of the mass-action system ($N\bar{x}$ with $x$ given by formula (4) or (8) in the case of the division-controlled or death controlled growth, respectively), then the above results do not hold, and the number of mutants in both fragmented and non-fragmented system is simply $N\bar{y}$, formula (5). Let us instead assume that the total population size, $N_{tot}$, is well below maximum. As a consequence, at size $N_{tot}$, each of the identical, disconnected populations in figure S7(b) that represent the mass action system, have not reached its maximum size. This is shown by a vertical dashed line that cuts across the growth phases of the $N$ patches in panel (b). A different picture is observed in the case of a fragmented system (panel (a)). There, since the growth in different patches happens at different times, by the time some of the patches have reached their maximum size, others have hardly started growing. As a consequence, at total size $N_{tot}$, we expect that a number of patches are "full" and others are "empty", see panel (b).

Now, we can formulate the problem of maximization of the number of mutants as size $N_{tot}$ in the following way. We can make up size $N_{tot}$ out of individual (identical) patches of size $x$ (precisely, $N_{tot}/x$ patches). The total number of mutants is then given by

$$M_{tot}(x) = N_{tot}\frac{\mathcal{M}(x)}{x}, \tag{22}$$

where $\mathcal{M}(x)$ is the number of mutants in a single patch of size $x$. What value of $x$ maximizes the function $M_{tot}(x)$? The answer depends on the function $\mathcal{M}(x)$. If, for example, it is a convex function, then quantity (22) is a growing function of $x$, and is maximized by a smaller number of patches, each at its maximum size ($x = \bar{x}$). If $\mathcal{M}(x)$ is a concave function, then it $M_{tot}$ is a decreasing function of $x$, and we find a maximum number of mutants if all the $N$ patches contribute the smallest possible amount into the total.

Functions $\mathcal{M}(x)$ have different shapes for different mutant types, see figure S3(b). For "decreased divisions" mutants, it is always a convex function, and thus $M_{tot}$ is maximized by a fragmented system, where at total size $N_{tot}$, a number of patches are already at carrying capacity while other have hardly began to grow.

For "increased death" mutants, the situation is slightly more complex. We know that these mutants grow faster than the wild type at the initial stages of growth (figure S3(a), yellow line), but depending on the degree of disadvantage, this growth may result in a monotonically growing function $y(t)$ for small degrees of disadvantage, or in a function $y(t)$ that "overshoots", for larger degrees of disadvantage. In the latter cases, the function $\mathcal{M}(x)$ has the shape depicted in figure S3(b), yellow line. As a consequence, function (22) will have a larger value for small $x$ (corresponding to the mass-action system where many virtual identical patches contribute a small amount) than for large $x$ (corresponding to the fragmented system).

Figure S8 shows the fraction of mutants as a function of total population size for "decreased divisions" and "increased death" mutants. For "decreased divisions" mutants, this is an increasing function of the total population size, and therefore to increase the percentage of mutants, one needs to maximize population size. This corresponds to having fewer patches at maximum size.

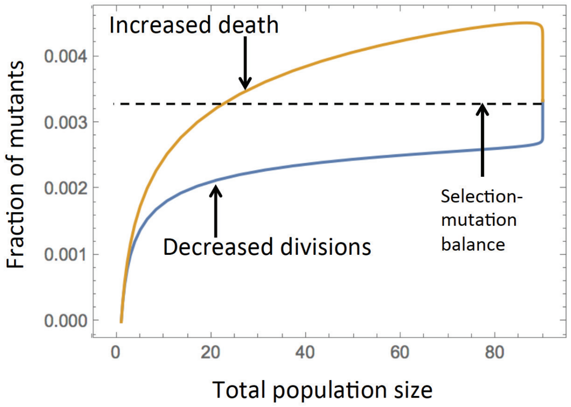

Figure S8: The fraction of mutants as a function of total population size for "decreased divisions" and "increased death" mutants. Parameters are as in figure S3.

For "increased death" mutants, the fraction of mutants first increases and the decreases. It is larger at an intermediate colony size compared to the maximum size for a large range of sizes, except for the initial stage of growth. Therefore, in most situations, a colony that is still growing will contain a larger percentage of mutants than the colony at maximum size. Therefore, we expect more mutations in the mass action situation where the saturation has not happened yet, and not in a fragmented system, where most colonies are at maximum size.

#### 2.7 Migrations on a complete graph model: deterministic and stochastic cases

Note that the intuitive explanation presented above is not specific for a 1D geometry and holds for any fragmented system where patches are de-synchronized, that is, they grow to their maximum size at different times. This is a necessary condition to be able to identify the stage of growth where a subset of patches is fully grown while the rest are empty. This is why the results described above (that is, a difference in the behavior of decreased divisions mutants and increased death mutants) is observed in stochastic simulations where no specific spatial arrangement of patches is assumed, and migration happens randomly from a patch to any other patch. In such a stochastic system, some patches will grow faster than others, and the general growth pattern similar to figure S7(a) is observed.

This behavior however is not captured by the corresponding system of ODEs. There, in the presence of equal migration rates to any patch, all patches (except for the original patch containing the first cell) are synchronized and grow in an identical manner. In fact, the ODEs in this case can be rewritten as only 2 equations in 2 patches:

$$\begin{aligned}
\dot{x}_* &= L_x x_*(1-u)\left(1-\frac{x_*+y_*}{K}\right) - D_x x_* - \mu\left(x_* - \frac{1}{N-1}X_*\right),\\
\dot{y}_* &= (L_x x_* u + L_y y_*)\left(1-\frac{x_*+y_*}{K}\right) - D_y y_* - \mu\left(y_* - \frac{1}{N-1}Y_*\right),\\
\dot{X}_* &= L_x X_*(1-u)\left(1-\frac{X_*+Y_*}{K(N-1)}\right) - D_x X_* - \mu\left(\frac{1}{N-1}X_* - x_*\right),\\
\dot{Y}_* &= (L_x X_* u + L_y Y_*)\left(1-\frac{X_*+Y_*}{K(N-1)}\right) - D_y Y_* - \mu\left(\frac{1}{N-1}Y_* - y_*\right),\\
& x_*(0)=1, \quad y_*(0)=0, \quad X_*(0)=0, \quad Y_*(0)=0.
\end{aligned}$$

This system can be thought of as a two-patch model, where the first patch has population $(x_*, y_*)$ and the carrying capacity $K$, and the second patch population $(X_*, Y_*)$ and the carrying capacity $K(N-1)$. The migration rate from the small to the large patch is $\mu$, and back it is $\mu/(N-1)$. Therefore, because of this artificial symmetry arising from the deterministic description (which is broken in a stochastic model), the behavior of this system does not reflect the patterns described above.

### 3 Disadvantageous mutants: a 2D spatial stochastic model

If mutants are disadvantageous, the quasi-equilibrium level of mutants is defined by the selection-mutation balance.

Let us calculate equilibrium densities of mutants and wild type cells in a spatially distributed system at steady state. This will also correspond to the densities in the core of an expanding system away from the advancing front.

We restrict our description to a 2D square grid, with the von Neumann neighborhood (that is, each location has 4 nearest neighbors); the methodology is generalizable to the Moore neighborhood (8 neighbors). We use a method similar to that of [2]. Two random variables describe the state of the stochastic system at each spatial location, $x$: $\rho_x$ describes wild type cells, such that

$$\rho_x = \begin{cases} 1, & \text{if a wild type cell is at location } x,\\ 0, & \text{otherwise,}\end{cases}$$

and $\eta_x$ describes mutant cells, such that

$$\eta_x = \begin{cases} 1, & \text{if a mutant cell is at location } x,\\ 0, & \text{otherwise.}\end{cases}$$

Note that $\rho_x$ and $\eta_x$ cannot be equal to one simultaneously; an empty spot corresponds to $\rho_x = \eta_x = 0$. We assume that wild type cells have division and death rates $L_x$ and $D_x$, and mutant cells have division and death rates $L_y$ and $D_y$. Wild type cells mutate with probability $u$, and no back mutations are considered.

##### 3.1 Equations for the densities

Denote the expectation of $\rho_x$ and $\eta_x$ by

$$\langle \rho_x \rangle = \rho, \quad \langle \eta_x \rangle = \eta,$$

where we assumed that the expected values do not depend on spatial location, since we are interested in spatially homogeneous equilibrium solutions. We have

$$\dot{\rho} = \left\langle \frac{L_x}{N_b}(1-u)(1-\rho_x)(1-\eta_x)\sum_k \rho_x^{(k)} - D_x\rho_x \right\rangle, \tag{23}$$

$$\dot{\eta} = \left\langle \frac{1}{N_b}(1-\rho_x)(1-\eta_x)\sum_k \left(L_x u \rho_x^{(k)} + L_y \eta_x^{(k)}\right) - D_y\eta_x \right\rangle, \tag{24}$$

where the product $(1-\rho_x)(1-\eta_x)$ is nonzero only if location $x$ is empty, and the summation goes over all the neighbors of point $x$, which reproduce into location $x$ at rates $L_x/N_b$ and $L_y/N_b$ if they are wild type of mutant, respectively.

Let us consider the von Neumann neighborhood ($N_b = 4$). In the right hand side of equation (23), the terms is the summation have the form

$$\langle (1-\rho_x)(1-\eta_x)\rho_x^{(k)} \rangle = \langle \rho_x^{(k)} \rangle - \langle \rho_x\rho_x^{(k)} \rangle - \langle \rho_x^{(k)}\eta_x \rangle + \langle \rho_x^{(k)}\rho_x\eta_x \rangle = \rho - W - I, \tag{25}$$

and in equation (24) there are also terms of the form

$$\langle (1-\rho_x)(1-\eta_x)\eta_x^{(k)} \rangle = \langle \eta_x^{(k)} \rangle - \langle \rho_x\eta_x^{(k)} \rangle - \langle \eta_x^{(k)}\eta_x \rangle + \langle \eta_x^{(k)}\rho_x\eta_x \rangle = \eta - I - M. \tag{26}$$

In the expressions above, we have $\langle \rho_x\rho_x^{(k)}\eta_x^{(k)} \rangle = 0$, because either $\eta_x^{(k)}$ or $\rho_x^{(k)}$ is zero at location $x^{(k)}$, and the three types of dyads are defined as follows:

- $W = \langle \rho_x\rho_x^{(k)} \rangle$ is the probability to have two wild type cells at two neighboring locations,
- $I = \langle \rho_x\eta_x^{(k)} \rangle$ is the probability to have a wild type cell and a mutant at two neighboring locations,
- $M = \langle \eta_x\eta_x^{(k)} \rangle$ is the probability to have two mutant cells at two neighboring locations.

Figure S9(a) illustrates these three configurations. In terms of these three correlations, equations (23) and (24) can be rewritten as

$$\dot{\rho} = L_x(1-u)(\rho - W - I) - D_x\rho, \tag{27}$$

$$\dot{\eta} = L_x u(\rho - W - I) + L_y(\eta - I - M) - D_y\eta. \tag{28}$$

The correlations for the three dyads that appear in these equations require their own equations to close the system. Let us derive an equation for $W$. We have

$$\dot{W} = \left\langle 2(1-\rho_x)(1-\eta_x)\rho_x^{(k)}\sum_j \rho_x^{(j)}\frac{L_x}{N_b}(1-u) - 2D_xW \right\rangle,$$

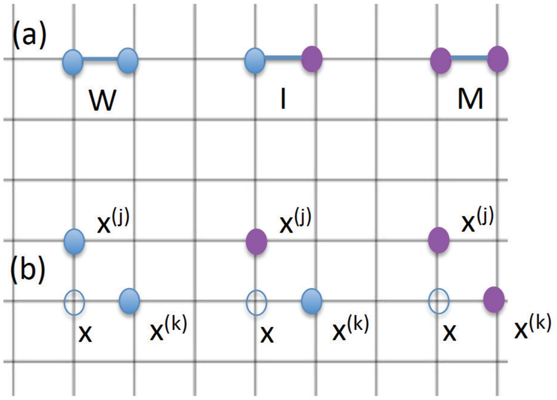

Figure S9: Steps in the derivation of equations for a two-component system of wild type and mutant cells. Blue circles denote wild type, and purple denote mutant cells. (a) Three configurations, whose correlations appear in equations (27) and (28). (b) Three types of correlations needed for equations for $W$, $I$, and $M$.

where we assume that one of the points in the dyad contains a wild type cell (term $\rho_x^{(k)}$), while the other point is empty (term $(1-\rho_x)(1-\eta_x)$), and that one of its neighbors (location $x^{(j)}$) contains a wild type cell, which reproduces faithfully into point $x$ at rate $L_x(1-u)/N_b$. Note that either of the two points could be empty, which results in the multiplier 2 in the first term on the right hand side. Similarly, either of the dyad's locations can experience cell death, resulting in the negative rate $2D_x$. In order to calculate the average, we need to consider terms

$$\langle (1-\rho_x)(1-\eta_x)\rho_x^{(k)}\rho_x^{(j)}\rangle. \tag{29}$$

Note that here and below, the operation of averaging makes the expression independent on the actual location $x$. Further, the superscripts $(k)$ and $(j)$ do not refer to any specific neighbor of $x$, but to any neighbor of $x$; in particular, location $x^{(j)}$ may be the same or different than location $x^{(k)}$. In the case when the two locations are different, correlation (29) is presented in figure S9(b), on the left.

In equations for $\dot{M}$ and $\dot{I}$, the following expressions appear in addition to (29):

$$\langle (1-\rho_x)(1-\eta_x)\rho_x^{(k)}\eta_x^{(j)}\rangle, \quad \langle (1-\rho_x)(1-\eta_x)\eta_x^{(k)}\eta_x^{(j)}\rangle.$$

These correlations are shown in figure S9(b), center and right. Therefore, denoting by $a$ and $b$ either $\rho$ or $\eta$, we evaluate the average of the form

$$\langle (1-\rho_x)(1-\eta_x)a_x^{(k)}b_x^{(j)}\rangle, \tag{30}$$

which corresponds to a dyad with one of the locations (location $x$) empty, and the other (location $x^{(k)}$) containing type "a", while a neighbor of $x$ (location $x^{(j)}$) contains type "b". First let us assume that location $x^{(j)}$ is different from location $x^{(k)}$. Under von Neumann neighborhoods this implies that $x^{(j)}$ are $x^{(k)}$ are not each other's neighbors, because on a square grid, there could not be a non-degenerate triangle with diameter 1 or less. Expression (30) is equal to

$$\begin{aligned} &P(b_x^{(j)} = 1 | a_x^{(k)} = 1, \rho_x = \eta_x = 0) P(a_x^{(k)} = 1, \rho_x = \eta_x = 0) \approx \\ &P(b_x^{(j)} = 1 | \rho_x = \eta_x = 0) P(a_x^{(k)} = 1, \rho_x = \eta_x = 0) = \\ &\frac{P(b_x^{(j)} = 1, \rho_x = \eta_x = 0) P(a_x^{(k)} = 1, \rho_x = \eta_x = 0)}{P(\rho_x = \eta_x = 0)} \end{aligned} \tag{31}$$

The expression in the denominator is calculated as follows:

$$P(\rho_x = \eta_x = 0) = \langle (1-\rho_x)(1-\eta_x) \rangle = \langle 1 - \rho_x - \eta_x + \rho_x \eta_x \rangle = 1 - \rho - \eta.$$

Depending on the types at location $x$, the expressions in the numerator of (31) can be of two types:

$$P(\rho_x^{(k)} = 1, \rho_x = \eta_x = 0) \text{ or } P(\eta_x^{(k)} = 1, \rho_x = \eta_x = 0),$$

and they are calculated in (25) and (26) respectively.

Next, we assume that location $x^{(j)}$ is the same as $x^{(k)}$. Then, if types "a" and "b" in expression (30) are different, then we obtain $\langle (1-\rho_x)(1-\eta_x)\rho_x^{(k)}\eta_x^{(k)} = 0$. If the types are the same, then we obtain expression (25) or (26). To summarize, expressions of type (30) are given as follows:

$$\begin{aligned} \langle (1-\rho_x)(1-\eta_x)\rho_x^{(k)}\rho_x^{(j)} \rangle &= \begin{cases} \frac{(\rho - W - I)^2}{1-\rho-\eta}, & x^{(j)} \neq x^{(k)}, \\ \rho - W - I, & x^{(j)} = x^{(k)}, \end{cases} \\ \langle (1-\rho_x)(1-\eta_x)\rho_x^{(k)}\eta_x^{(j)} \rangle &= \begin{cases} \frac{(\rho - W - I)(\eta - I - M)}{1-\rho-\eta}, & x^{(j)} \neq x^{(k)}, \\ 0, & x^{(j)} = x^{(k)}, \end{cases} \\ \langle (1-\rho_x)(1-\eta_x)\eta_x^{(k)}\eta_x^{(j)} \rangle &= \begin{cases} \frac{(\eta - I - M)^2}{1-\rho-\eta}, & x^{(j)} \neq x^{(k)}, \\ \eta - I - M, & x^{(j)} = x^{(k)}. \end{cases} \end{aligned}$$

The equation for $W$ is then given by

$$\dot{W} = \frac{L_x}{2}(1-u)\left(\rho - W - I + \frac{3(\rho - W - I)^2}{1-\rho-\eta}\right) - 2D_x W. \tag{32}$$

Similarly, the other two equation can be derived:

$$\begin{aligned} \dot{I} &= \frac{3}{4}[L_x(1-u) + L_y]\frac{(\rho - W - I)(\eta - I - M)}{1-\rho-\eta} + \frac{L_x u}{4} L_y \left(\rho - W - I + \frac{3(\rho - W - I)^2}{1-\rho-\eta}\right) \\ &- (D_x + D_y) I, \end{aligned} \tag{33}$$

$$\dot{M} = \frac{3L_x u}{2}\frac{(\rho - W - I)(\eta - I - M)}{1-\rho-\eta} + \frac{L_y}{2}\left(\eta - I - M + \frac{3(\eta - I - M)^2}{1-\rho-\eta}\right) - 2D_y M. \tag{34}$$

The closed system of equations for $\rho, \eta, W, I$, and $M$ is given by equations (27), (28), (32), (33), and (34).

##### 3.2 Selection mutation balance solution

Solving these equations in steady state exactly is difficult, but if the mutation rate $u \ll 1$, we can find the approximate solution. We start by setting $u = 0$ and obtaining the steady state solution. Apart from the trivial solution and a negative solution, there are two symmetric solutions where only one type survives (competitive exclusion). We will use the one where the wild type excludes mutants:

$$\rho^{(0)} = 1 + \frac{3D_x}{D_x - 3L_x}, \quad W^{(0)} = 1 - \frac{6D_x}{D_x - 3L_x} - \frac{4D_x}{L_x}, \quad \eta^{(0)} = I^{(0)} = M^{(0)} = 0, \tag{35}$$

where the superscript corresponds to the zeroth order in the expansion in terms of small $u$. Note that the expression for $\rho^{(0)}$ is identical to the steady state density of the one-component system given by equation (**??**) with $\xi = D_x/L_x$. We then look for the first correction by substituting

$$\rho = \rho^{(0)} + u\rho^{(1)}, \quad \eta = u\eta^{(1)}, \quad W = W^{(0)} + uW^{(1)}, \quad I = uI^{(1)}, \quad M = uM^{(1)},$$

inserting in the system of 5 equations, keeping only the first order of expansion in $u$, and solving for $\rho^{(1)}, \ldots, M^{(1)}$. We obtain

$$\eta = \eta^{(1)}u = \frac{D_x L_x (4D_x - 3L_x)(4D_y^2 + 3D_y L_x + D_x L_y)u}{D_y (D_x - 3L_x)(4D_y + 3L_x)(D_y L_x - D_x L_y)}.$$

This is the equilibrium solution corresponding to mutation-selection balance in the presence of spatial interactions. This approach is valid as long as the wild type is advantageous (inequality (3)). In the opposite scenario, this solution is unstable, and the system converges to the mutants excluding the wild type.

Under selection-mutation balance, of interest is the equilibrium proportion of mutants in the system given by

$$\nu_{vN} = \frac{\eta^{(1)}u}{\rho^{(0)}} = \frac{D_x L_x (4D_y^2 + 3D_y L_x + D_x L_y)u}{D_y (4D_y + 3L_x)(D_y L_x - D_x L_y)}. \tag{36}$$

In the absence of death ($D_x = D_y = 0$), we obtain the limiting value

$$\nu_{vN,D=0} = \frac{(3L_x + L_y)u}{3(L_x - L_y)}.$$

##### 3.3 Comparison with mass-action

We would like to compare the proportion of mutants under selection mutation balance in space and in mass-action. In mass action we have

$$\dot{x} = L_x(1-u) - D_x x, \tag{37}$$

$$\dot{y} = = L_x u x + (L_y - D_y) y, \tag{38}$$

where $x$ and $y$ is the expected number of wild type and mutant cells respectively. The proportion of mutant cells at a given size $N$ is calculated by using $x^{(0)} = e^{(L_x - D_x)t}$ and evaluating the solution $y(t)$ of equation (38) at time $t = \ln N/(L_x - D_x)$. Forming the fraction, we obtain the proportion at mutants at size $N$:

$$\nu_{ma} = \frac{L_x u \left(1 - N^{-\frac{L_x - D_x - (L_y - D_y)}{L_x - D_x}}\right)}{L_x - D_x - (L_y - D_y)}.$$

One can see that this quantity grows with the size $N$, while $\nu_{vN}$ is independent of the population size.

In figure S10 we study the quantity

$$\ln\left(\frac{\nu_{vN}}{\nu_{ma}}\right),$$

which is greater than 0 of the numerator is larger than the denominator. This quantity is presented as a contour plot in panel (a), as we vary $L_y$ and $D_y$ (for fixed values of $L_x$ and $D_x$). The red line in the figure indicates the region where the mutants are disadvantageous and we expect to observe selection-mutation balance (this is given by inequality (3)). Positive regions (below the purple dashed line) correspond to having more mutants in the spatial system compared to the mass action system. Negative regions (above the purple dashed line) are regions where there are fewer mutants in the spatial system than in the mass action system.

Figure S10(b) focuses on two particular scenarios. The top panel presents the case where the division rates of mutants and wild type cells are the same, and the disadvantage is manifested through $L_y < L_x$. We can see that in such cases, $\nu_{vN} > \nu_{ma}$, that is, we have more mutants in space. The bottom panel shows the case where $L_y = L_x$, and the mutants have a larger death rate than the wild types. In this case, if the disadvantage is sufficiently large, we have fewer mutants in space than in the mass action system.

#### 3.4 Comparison with computations

The expected number of mutants predicted theoretically was compared with results of numerical simulations. This was done in the following way. At size $N$, the number of mutants (in the von Neumann case) is predicted to be $Nu\nu_{vN}$, see equation (36). Solving the equation $Nu\nu_{vN} = const$, we can obtain the pairs $(L_y, D_y)$ of mutant kinetic rates corresponding to a predicted given number of mutants in a system of size $N$. Figure S11(a) shows the predicted number of mutant as a contour plot. The closer to the "neutrality" line (see inequality (3)) the larger the predicted number of mutants. Solution of equation $Nu\nu_{vN} = const$ is shown in panel (b), and for 5 points from the solution set, the predicted number of mutants (given by 10) is compared with the numerically obtained mean (plotted together with the standard deviation, panel (c)). We can see that for larger mutant division rates, the deviation from the theory becomes significant. Panel

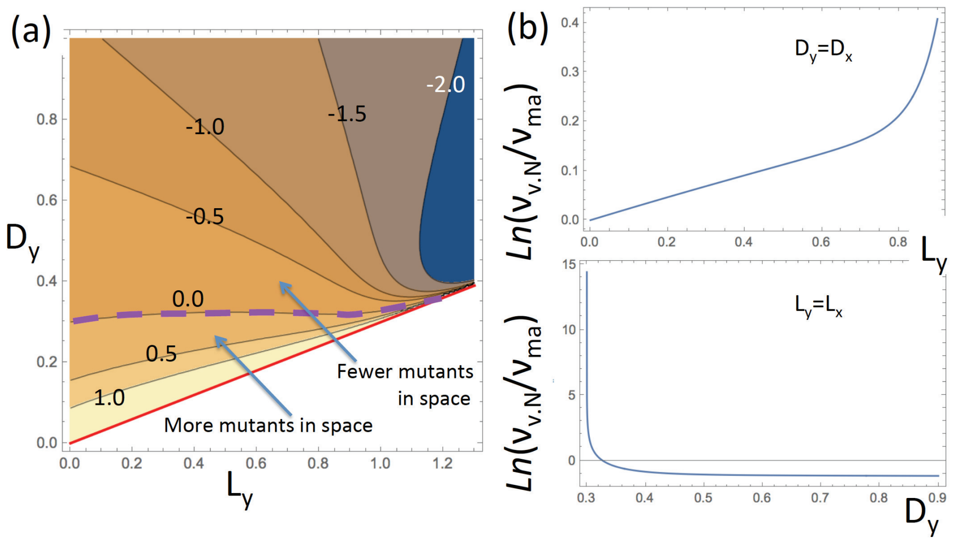

Figure S10: Comparison of the spatial and mass action system. (a) The quantity $\ln\left(\frac{\nu_{vN}}{\nu_{ma}}\right)$ is presented as a contour plot as a function of $L_y$ and $D_y$, for fixed values of $L_x$ and $D_x$. The red line indicates the region where mutants are disadvantageous (inequality (3)). The contours's values are indicated, and the zero contour is marked with a dashed purple line. (b) Top panel: the same quantity plotted as a function of $L_y$, with $D_y = D_x$. Bottom panel: the same, as a function of $D_y$, with $L_y = L_x$. The rest of the parameters are given by $u = 2 \times 10^{-5}$, $N = 10^5$, $L_x = 0.1$, and $D_x = 0.3$.

(d) shows a histogram of numbers of mutants for the parameters corresponding the 5th point. One can see that the distribution has a long tail and a very large standard deviation. This is the consequence of "slices" that cannot be handled by the present method.

#### 3.5 Jack-pot events

In Fig. 2 of the main text we studied the mean and the distribution of the number of disadvantageous mutants in mass-action systems and in a metapopulation model, were the demes were arranged in a 1D array, and migration happened between neighboring demes. Here we present results for a metapopulation model where migration was equally likely among all demes, see figure S12. We can see that regardless of the structure of the deme-to-deme network, the results are very similar.

In order to compare the number of rare ("jackpot") events in different models and for different parameters, we designed a function that quantifies the spread

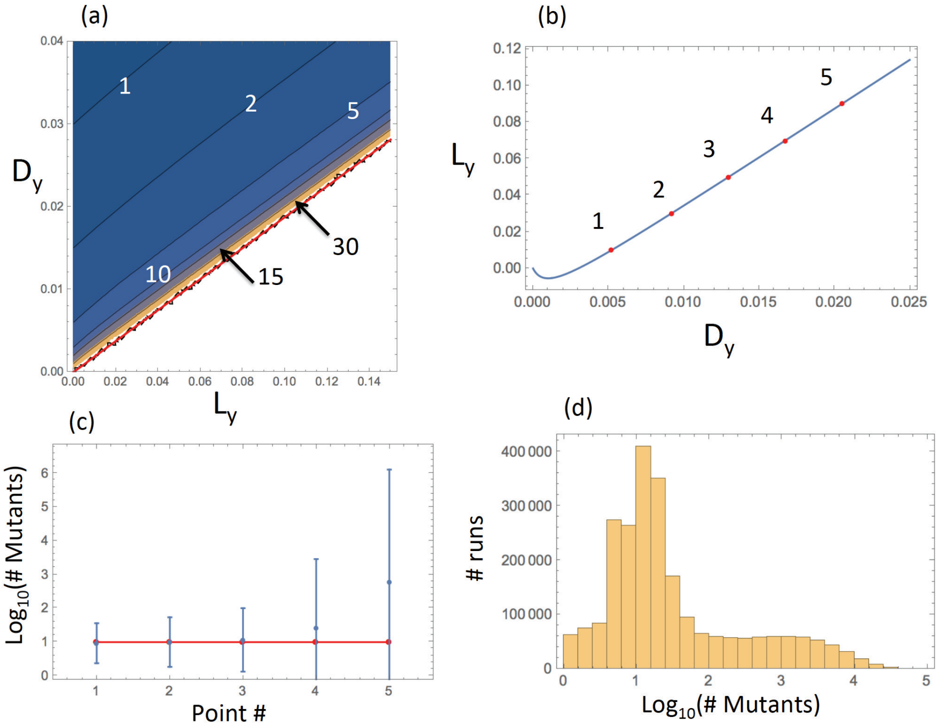

Figure S11: The level of mutants in the spatial (von Neumann) system: analytical approximation and numerical results. (a) The quantity $Nu\nu_{vN}$ is presented as a contour plot as a function of $L_y$ and $D_y$, for fixed values of $L_x$ and $D_x$. Mutants are disadvantageous above the red line (inequality (3)). The contours' values are specified. (b) Solution $L_y$ of equation $Nu\nu_{vN} = 10$ as a function of $D_y$; the 5 points used in panel (c) are marked in red and numbered. (c) The comparison of predicted (10, horizontal red line) and simulated number of mutants in the 5 parameter pairs from panel (b). Simulated means and standard deviations are shown (out of $2.5 \times 10^6$ runs). (d) For the 5th parameter combination, the numerically obtained histogram of the number of mutants is shown. The rest of the parameters are $u = 2 \times 10^{-5}$, $N = 10^5$, $L_x = 0.08$, and $D_x = 0.15$.

of the distribution or the "fatness" of its tail (heavy-tailedness). For each set of data, $Y$, representing the numbers of mutants in each of the runs performed for a given parameter set for a given model, we used the quantile function

$$F(Y, q) = Q(Y, 1 - q),$$

defined as follows. Let $Y_{qL}$ and $Y_{qR}$ are two subsets of $Y$ such that $Y = Y_{qL} \cup Y_{qR}$ and all the elements of $Y_{qL}$ are smaller or equal to all the elements of $Y_{qR}$; we

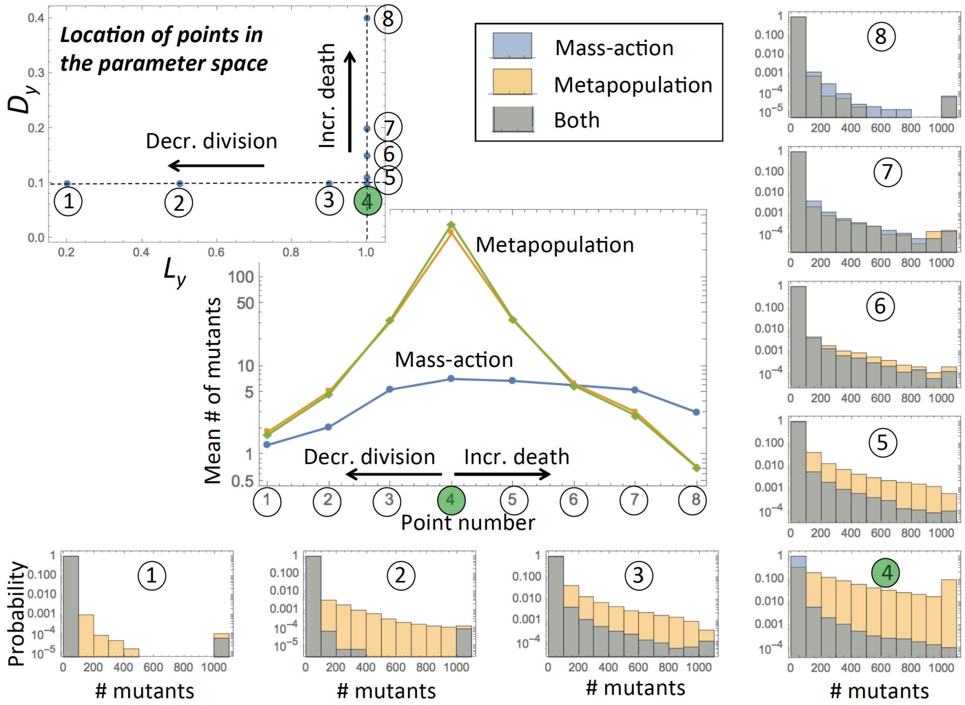

Figure S12: Disadvantageous mutants: probability distributions of mutant numbers in mass-action and metapopulation simulations. Both decreased divisions and increased death mutants are investigated, with division and death rates given in the left upper panel (compared to the rates of the wild type, denoted by the green circle). The bar graphs represent numerical histograms for mass action simulations (blue) and metapopulation simulations (yellow), where the demes were all connected to each other; between 180000 and 200000 simulations were run for each parameter combination, and the number of mutants recorded when the total population reached 1000. The mean numbers of mutants are shown for all the simulations in the central graph, with green line corresponding to the present metapopulation model, and the yellow line to the 1D metapopulation model with only near neighbor connection, presented in Fig. 2 of the main text.

will refer to parameter $q$ as "threshold". We further assume that

$$|Y_{qR}| = \lceil |Y|q \rceil,$$

where $|.|$ denotes the number of elements and $\lceil . \rceil$ denotes ceiling. Then we set

$$Q(Y, 1-q) = \min(Y_{qR}).$$

It follows from this definition that the function $F(Y_1, q) > F(Y_2, q)$ for all $q$ in a vicinity of 0, as long as set $Y_1$ has a higher number of large outliers compared to $Y_2$, or if $Y_1$ is drawn from a distribution with a heavier tail compared to $Y_2$.

Figure S13 presents the plots of the heavy-tailedness function $F(Y, q)$ for the 8 cases studied in figure S12, with $Y$ representing the mass-action model (blue), the metapopulation model where all demes are connected (yellow), and the 1D metapopulation model with only neighboring demes connected (green). The bottom row of graphs represents the cases where the disadvantageous mutants are characterized by a decreased division rate (except case 4, which describes neutral mutants). In the top row the disadvantageous mutants are characterized by increased death rates. We can see that the heavy-tailedness generally increases toward the neutral case. We also observe the following patterns:

- In the bottom row, for small thresholds, $q$, the yellow and green lines are above the blue line, which means that if the mutants have decreased division rates, the fragmented (metapopulation) models are characterized by a higher heavy-tailedness (more jack-pot events) compared to the well-mixed system.

- This trend weakens and reverses in the graphs of the top row, from right to left (away from neutral mutants). In other words, if the mutants have increased death rates, and the disadvantage is sufficiently large, the fragmented (metapopulation) models are characterized by a lower heavy-tailedness (fewer jack-pot events) compared to the well-mixed system.

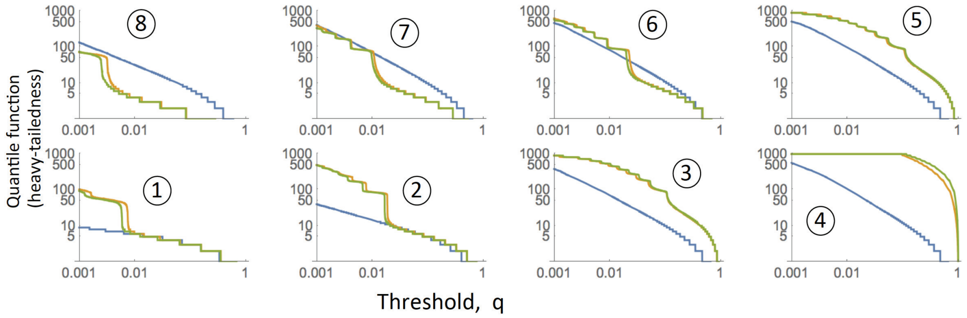

Figure S13: The prevalence of jack-pot events in different models. The heavy-tailedness (quantile) function $F(Y, q)$ is plotted vs $q$ for the mass-action model (blue), the metapopulation model where all demes are connected (yellow), and the 1D metapopulation model with only neighboring demes connected (green). The calculations are based on the simulations of figure S12 and figure 2 of the main text. Case numbering and parameters are as in figure S12.

These trends go hand in hand with the results on the expected number of mutants (figure S12, middle graph). For mutants with decreased division rates or slightly increased death rates, there are more jack-pot events in the metapopulation models, and the expected number of mutants is also higher, compared to the well-mixed model. On the other hand, if the mutants are characterized by significantly higher death rates, the trend reverses, and there are more jack-pot events and more mutants on average in the mass-action system.

### 4 Neutral and advantageous mutants in a range expansion

#### 4.1 Derivation of the growth laws

To derive the laws of mutant growth reported in the main text, we can use the following simple calculations. Let us assume that the death rate of cells is equal to zero, and consider cells growing in different geometries.

**1D: surface of a cylinder.** Assume that cells grow along the surface of a cylinder of width $W$. This represents a one-dimensional growth process, where during each generation, we assume that $W$ new cells appear, and the the total population is given by $N = LW$, where $L$ represents the number of layers. The value of $L$ is proportional to the number of generations, and thus to the physical time, $t$:

$$L \propto t.$$

The following calculation estimates the growth law of mutants. Every time a new layer (of width $W$) is added, the mean number of new mutations is given by $Wu$. Suppose that mutants are neutral. Then, each such mutation will give rise to an array of daughter mutant cells of width 1, see figure S14. The length of this array is given by $L - i$, if $i$ is the layer at which the mutation occurred. Therefore, the total expected number of neutral mutants is a cylinder of length $L$ is given by

$$M_{1D}^{neut} = \sum_{i=1}^{L} Wu \sum_{j=i+1}^{L} 1 = \frac{uWL(L-1)}{2} \approx \frac{uWL^2}{2} = \frac{u}{2W}N^2, \tag{39}$$

where we assumed $L \gg 1$. Note that in this derivation we assumed that the number of mutants is small compared to the total population, and individual mutant clones do not interact. In a more precise calculation, the number of wild type cells in each layer is smaller than $W$ because of the existence of mutants, and thus the ate of new mutant production is smaller than $Wu$. We however assume that $uLW \ll 1$, such that the number of mutants is relatively small.

Note that the number of neutral mutants decreases with $W$, see figure S15; the largest number of mutants is achieved in the case of $W = 1$, a one-dimensional expanding array of cells.

Next, let us consider advantageous mutants. In this case, each new mutant gives rise to a triangular clone. In the first layer, the width of the clone is 1, in

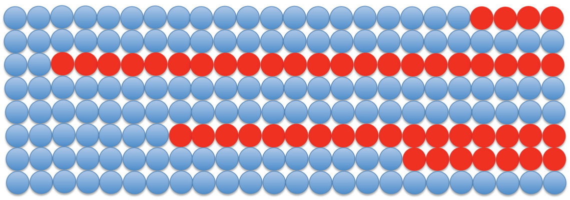

Figure S14: The conceptual model for mutant number calculations, the case of neutral mutants in a colony growing along the surface of a 1D cylinder.

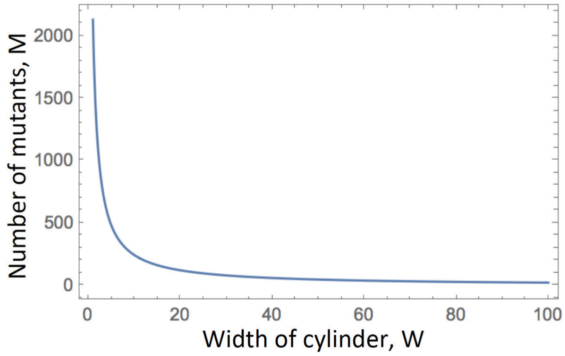

Figure S15: The number of mutants in a 1D cylinder decays with its width. Formula (39) is presented with $N = 10000$ and $u = 5 \times 10^{(-5)}$.

the next layer it is $1+s$, and in the $k$th layer it is $1+(k-1)s$, where parameter $s \geq 0$ measures the advantage of the mutants (with $s = 0$ corresponding to neutral mutants). Therefore, we have

$$M_{1D}^{adv} = \sum_{i=1}^{L} Wu \sum_{j=i+1}^{L} (1+(j-(i+1))s) = uWL(L-1)\left(\frac{1}{2}+\frac{s(L-2)}{6}\right) \qquad (40)$$
$$\approx \frac{uWsL^3}{6} = \frac{us}{6W^2}N^3,$$

where for the approximation, we assumed that $Ls \gg 1$. Also, for this simple calculation to be valid, we need to assume that the wedges created by mutants do not come close to the cylinder's width, $W$, that is, $Ls \ll W$. In particular, formula (40) can be valid for small values of $W$ and even $W = 1$, but only for mutants that are neutral for practical purposes ($s \ll 1$).

Note that in the special case where $W = 1$, mutant advantage does not play a role, and the number of advantageous, neutral, and even disadvantageous mutants is given by the same formula, equation (39).

**2D: circular range expansion.** Next we turn to the dynamics of neutral mutants on a circle. Let us suppose that the radius of the circle is $R$ and $N = \pi R^2$. The size of the colony increases via surface growth with $N \propto t^2$ and

$$R \propto t.$$

As the range expansion proceeds, the circular layer of radius $r$ will on average give rise to $2\pi r u$ new mutations. Each mutation will result in a wedge expanding outwards. If the new mutation occurred in layer with radius $r$, the number of mutating cells in layer $r$ is 1. The number of mutants in the next layer is given by $\frac{r+1}{r}$, because under the assumption of mutant neutrality, the fraction of mutants in each new layer of radius $j > r$ (with surface $2\pi j$) should stay constant and equal to $\frac{1}{2\pi r}$. For layer $j$, the number of mutants is then given by $j/r$. This gives rise to the following calculation:

$$M_{2D}^{neut} = \sum_{r=1}^{R} 2\pi r u \sum_{j=r+1}^{R} \frac{j}{r} = \frac{2}{3}\pi R(R^2-1)u \approx \frac{2\pi R^3 u}{3} = \frac{2u}{3\pi^{1/2}} N^{3/2}$$

(the approximation is valid for $R \gg 1$).

For advantageous mutants in a growing 2D circle, the fraction of mutants will grow with each layer:

$$\begin{aligned} M_{2D}^{adv} &= \sum_{r=1}^{R} 2\pi r u \sum_{j=r+1}^{R} (1+(j-(r+1))s)\frac{j}{r} = \pi R(R^2-1)u\left(\frac{2}{3}+\frac{1}{4}s(R-2)\right) \\ &\approx \frac{\pi R^4 s u}{4} = \frac{su}{4\pi} N^2, \end{aligned} \tag{41}$$

where we assumed $Rs \gg 1$. For this approximation to be valid, the mutant wedges should not exceed the circumference of the colony. Strictly speaking, this results in the condition $Rs << 2\pi R$, that is, $s \ll 1$. For larger values of $s$, the events where the mutant covers the whole surface of the colony are no longer negligible.

**3D: solid cylinder growing lengthwise.** In a 3D space, let us first consider a solid cylinder of constant radius $R_0$, where initially the cells are situated as a layer at the bottom of the cylinder, and proceed to grow by adding layers of size $\pi R_0^2$. Each generation contributes $\pi R_0^2 u$ new mutants, and as the colony grows to length $L$ (and volume $2\pi R_0^2 L$), we have in the neutral case:

$$M_{solid\ cyl}^{neut} = \sum_{i=1}^{L} 2\pi R_0^2 L \sum_{j=i+1}^{L} 1 = \pi R_0^2 u L(L-1) \approx \pi R_0^2 u L^2 = \frac{u}{\pi R_0^2} N^2,$$

which is similar to the 1D growth. If the mutants are advantageous, then their number will increase from layer to layer, giving rise to conical wedges. This gives rise to the following calculation:

$$
\begin{aligned}
M^{adv}_{solid\ cyl} &= \sum_{i=1}^{L} 2\pi R_0^2 L \sum_{j=i+1}^{L} (1+(j-(r+1))s)^2 \\
&= \frac{L(l-1)\pi R_0^2 u}{6}\left[(L^2-3L+2)s^2+4(L-2)s+6\right] \\
&\approx \frac{\pi R_0^2 s^2 u L^4}{6} = \frac{s^2 u}{6\pi^3 R_0^6} N^4,
\end{aligned}
$$

where $Ls \gg 1$ for the approximation, and the approach is valid as long as the wedge radius is smaller than that of the cylinder, $sL \ll R_0$.

**3D: spherical range expansion.** Next we consider a 3D expanding sphere. For a sphere of radius $R$, we have $N = 4/3\pi R^3$ and the surface is given by $4\pi R^2$. The size of the colony increases via 3D surface growth with $N \propto t^3$. Each spherical layer of radius $r$ will on average give rise to $4\pi r^2 u$ new mutations. Each mutation will result in a conical wedge expanding outwards. If the new mutation occurred in layer with radius $r$, the number of mutating cells in layer $r$ is 1. The number of mutants in a layer of radius $j > r$ is given by $(j/r)^2$, because under the assumption of mutant neutrality, the fraction of mutants in each new layer should stay constant (and equal to $\frac{1}{4\pi r^2}$). Therefore, we write:

$$
M^{neut}_{3D} = \sum_{r=1}^{R} 4\pi r^2 u \sum_{j=r+1}^{R} \left(\frac{j}{r}\right)^2 = \pi R(R^2-1)(R+2/3)u \approx \pi R^4 u = \frac{3^{4/3}u}{\pi^{1/3}4^{4/3}} N^{4/3}
$$

(the approximation is again valid for $R \gg 1$).

If the mutant in a growing 3D sphere is advantageous, the fraction in each layer will increase according to the fitness advantage $s$ and stretch from layer to layer in the same way as for the neutral mutants. We therefore have,

$$
\begin{aligned}
M^{adv}_{3D} &= \sum_{r=1}^{R} 4\pi r^2 u \sum_{j=r+1}^{R} (1+(j-(r+1))s)^2 \left(\frac{j}{r}\right)^2 \\
&= \frac{\pi u R(R^2-1)}{90}\left[(20R^3-48R^2-5R+42)s^2+(72R^2-90R-108)s+90R+60\right] \\
&\approx \frac{2}{9}\pi s^2 u R^6 = \frac{s^2 u}{8\pi} N^2.
\end{aligned}
$$

As before, the approximation holds if $Rs \gg 1$. The method assumes that the mutant colony's size in each layer does not come close to the surface area, which amounts to the inequality $s \ll 1$.

**Exponential (non-spatial, mass-action) growth.** Finally, for exponentially growing population, similar formulas could be derived. In particular, for neutral mutants, we have

$$
M^{neut}_{exp} = Nu \ln N,
$$

and for advantageous mutants with advantage $\alpha$ (which is the ratio of the net growth rate of mutants and the net growth rate of wild type cells), we have

$$M_{epx}^{adv} = \frac{\alpha}{(\alpha-1)2^{\frac{\alpha-1}{\alpha}}} N^{\frac{2\alpha-1}{\alpha}},$$

see [3], equation (14c), see also equation (13) for a more general formula.

A summary of some of the results is presented in table 1 of the main text.

#### 4.2 Comparison with numerical simulations

We have run numerical simulations to check the power laws for the neutral and advantageous mutants derived in the previous section.

##### 4.2.1 Growth on a cylinder (1D growth)

For the 1D (cylindrical) geometry, we set up the initial condition for the agent-based simulation to be a one layer (a circle) of wild type cells that coincides with the circumference of one of the cylinder's bases (and has length $W$). Simulations are run repeatedly for a fixed number of time steps, and the mean trajectory (that is, the mean number of wild type cells and the mean number of mutants, as functions of time-step) is then calculated. Finally, the number of mutants is plotted as a function of the total population size.

Examples for neutral mutants (that is, mutants that have the same division and death rates as the wild type cells) are presented in figure 4 of the main text, curves (a) and (b). We can see that both in the absence (a) and in the presence (b) of death, the mutant population as a function of the total population approaches a power law with the exponent 2 (the black dashed lines corresponding to cases (a) and (b) of figure 4 of the main text have slope 2 in the log-log plot, see table 1). There are more mutants in the presence (b) than in the absence (a) of death.

Advantageous mutants are presented by curves (c-g) of figure 4 of the main text; note that the dashed lines drawn through these curves all have slope 3 in the log-log coordinates (table 1). Curves (c) and (d) correspond to systems without death, and mutants having a larger division rates compared to wild type cells. The advantage is larger in case (d) compared to case (c) (and thus there are more mutants observed at the same population size). In both cases, we can see that the curves have slope 3 up to a certain population size, after which they deviate from the cubic law. For those larger sizes, mutants grow slower as a function of size (quadratically). The reason for this deviation from the cubic law is as follows. As the colony grows and reaches larger sizes, advantageous mutant clones that grow out and increase in size start reaching the width $W$, that is, take up the whole width of the cylinder. After that, the mutant colony can no longer expand in width, but instead it grows linearly. This is illustrated in figure S16(a), where the top panel replots the purple line (case (d)) of figure 4 of the main text, and the bottom panel studies the statistics of mutant invasion. For each run, we recorded the total colony size at which the

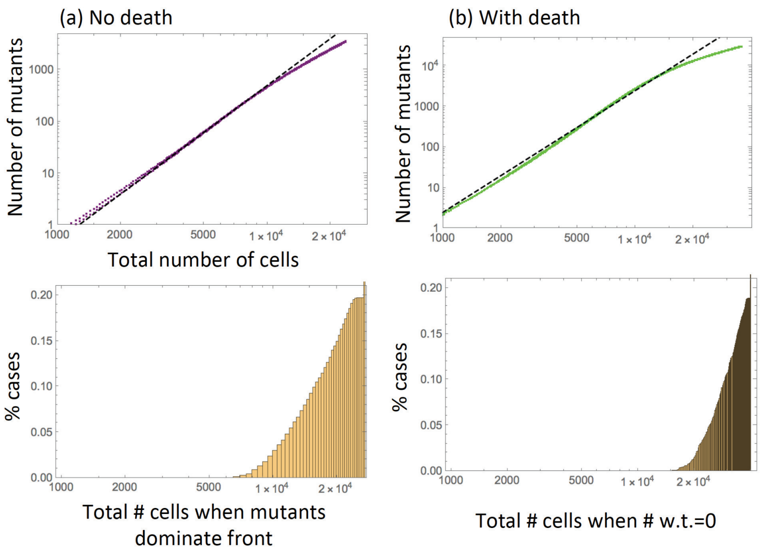

Figure S16: Advantageous mutants on a cylinder: deviation from the cubic law for large sizes. (a) In the absence of death ($L_x = 0.7, L_y = 1.0$, $D_x = D_y = 0$, see curve (d) of figure 4 of the main text) and (b) in the presence of death ($L_x = L_y = 0.7$, $D_x = 0.2, D_y = 0.1$, see curve (f) of figure 4). Top panels: the number of mutants as a function of the total number of cells; the dashed straight lines have slope 3. Bottom panels: (a) a numerically obtained CDF of the probability for a given colony to have its front completely dominated by mutants by a given size; (b) a numerically obtained CDF of the probability for a given colony to have its wild type population extinct by a given size.

population of wild type cells stopped increasing (given that this happened in the time-span of the simulation). This indicates that, in the absence of death, all the front positions are taken up by mutants and wild type cells can no longer divide. The numerically obtained CDF is presented in the panel. We can see that the probability for the mutants to dominate the front becomes significant around size $10^4$, which coincides with the size where the number of mutants starts deviating from the cubic law (the upper panel).

Next, we turn out attention to curves (e) and (f) of figure 4. They represent systems in the presence of death, where mutant advantage is manifested through increased division rate (e) and decreased death rate (f). We can see again that the curves follow a cubic law. A deviation from this law (and a slow-down of the growth as a function of total size) is also observed for larger sizes. The mechanism of this deviation is however somewhat different from the case of zero death. Figure S16(b, top panel) replots curve (f) of figure 4. Since cells die, the eventual outcome of all the simulations is the extinction of the (disadvantageous) wild type. The CDF of the colony size by which the wild type goes extinct is presented in the bottom panel of figure S16(b). The probability of wild type extinction becomes significant around size $2 \times 10^4$, where the number of mutants starts deviating from the cubic growth (see the top panel).

Finally, we compare curves (f) and (g) of figure 4. They represent systems with the same parameters (where the mutant advantage is manifested through a lowered death rate), except the cylinder width is $W = 100$ for curve (f) and it is $W = 1000$ for curve (g). Notice that the dashed black lines for the two curves differ by a factor of 100, representing the inverse square dependence of the number of mutants on the cylinder width, see formula (40) (there are 100 times fewer cells in the cylinder that is 10 times wider).

##### 4.2.2 Growth on a circle (2D growth)

For simulations studying 2D growth, we started with 1 wild cell, and let the colony expand on a 2D grid for 800 time-steps, for. 2000 runs for each parameter combination. In each case, the numbers of wild type and mutant cells were averaged over all the runs that did not result in population extinction[1]. The results for several representative cases are presented in figure 6 of the main text, where for each parameter combination the number of non-extinct runs is given in the figure caption.

There are 6 curves plotted in figure 5 of the main text. As in figure 4, cases (a,b) correspond to neutral mutants, and cases (c-f) to advantageous mutants; please note that all the division and death rates in curves (a-f) of figure 5 are identical to curves (a-f) of figure 4.

The black dashed lines for curves (a,b) in figure 5 have slope 3/2, as predicted for neutral mutant growth in a circle. As in the case of the cylinder, there are more mutants in the presence (b) than in the absence (a) of death.

The slope for curves (c-f) in figure 5 is 2, as predicted for the growth of advantageous mutants in a circle, see table 1. Again, we observe deviation fro the predicted power law for large system sizes. In the absence of cell death (cases (c,d)) this happens as the mutants become more likely to spread and occupy all the surface locations, blocking the wild type cells from divisions. In the presence of death, this deviation is associated with the increased likelihood of wild type extinction.

[2] Ignacio A. Rodriguez-Brenes, D. Wodarz, and N.L. Komarova. Beyond the

[1]Extinction was not a problem in the cylindrical geometry, because the initial condition contained 100 or more cells, and very few runs resulted in extinction

pair approximation: modeling colonization population dynamics. *In submission*, 2019.

[3] Yoh Iwasa, Martin A Nowak, and Franziska Michor. Evolution of resistance during clonal expansion. *Genetics*, 172(4):2557–2566, 2006.
